# Mechanical characterization of human centromeres

**DOI:** 10.64898/2026.10.14.759046

**Authors:** Tinka V.M. Clement, Christian F. Nielsen, Zhaowei Liu, Bram Hoogland, Rui Cui, Colleen C. Caldwell, Chase P. Broedersz, Ian D. Hickson, Gijs J.L. Wuite

## Abstract

Centromeres must withstand high pulling forces in metaphase, yet allow rapid and complete chromosome segregation in anaphase. Using optical tweezers measurements on isolated human metaphase chromosomes, we find that centromeres are stiffer at low forces and more frequently exhibit pronounced softening compared to whole chromosomes, while displaying a similar stiffening response at high forces. We capture these mechanical properties by establishing a softened hierarchical worm-like chain model. Cohesin cleavage alters the mechanical properties of centromeres but does not lead to chromosome segregation *in vitro*. Topoisomerase IIα is recruited to centromeric regions under significant tension, likely to resolve the persistent centromeric DNA entanglements that provide additional sister chromatid cohesion. These findings advance our understanding of the mechanical resilience and regulation of centromeres during mitosis.

---

Faithful chromosome segregation is essential for the successful progression of mitosis and for the prevention of genome instability in daughter cells. Centromeres are specialized chromosomal regions where the sister chromatids are linked together and connect to the mitotic spindle. Human regional centromeres consist of highly repetitive arrays of DNA in core centromeres flanked by divergent, less repetitive DNA arrays in pericentromeres(*1*, *2*). A subset of the nucleosomes in core centromere regions contain the histone H3 variant CENP-A(*3*, *4*). CENP-A-rich regions of core centromeres direct the formation of kinetochores, protein complexes that function as the attachment point for spindle microtubules and are responsible for force transduction between the mitotic spindle and centromeric DNA(*5*, *6*).

To ensure sister chromatid cohesion up to metaphase, cohesin, a ring-shaped protein complex that can encircle multiple strands of DNA(*7*, *8*), links DNA from both sister chromatids near the centromeres(*7–10*). *In vivo*, cleavage of the cohesin subunit SCC1 (RAD21) by separase triggers separation of sister chromatids and marks the onset of anaphase(*10–12*). In addition to cohesin cleavage, any remaining centromeric DNA entanglements between sister chromatids also need to be resolved. These entanglements occur in all mitoses and originate from dsDNA catenanes(*13–19*). They are present already in metaphase and their resolution at the metaphase-anaphase transition or during anaphase is essential for successful mitotic exit(*13*, *16*, *20–22*).

The structural integrity of centromeres must be maintained throughout mitosis, despite the mitotic spindle applying pulling forces of tens to hundreds of piconewtons(*23–26*). This raises questions about the specific mechanical properties that allow centromeres to carry out its functions. Studies on centromere mechanics have previously been performed using (super-resolution) microscopy combined with advanced image analysis and modelling(*27–29*). These methods provide an indirect means of controlling forces and determining centromere mechanics. Micropipette aspiration(*30*, *31*) and dual-trap optical tweezers(*32–34*) do allow for direct application of forces to isolated native mitotic chromosomes by stretching them. Optical tweezers measurements have been used for mechanistic modelling of mitotic chromosome organisation(*32*, *34*). Similar progress would be essential for understanding the mechanical properties of centromeres and the mechanisms involved in ensuring controlled and faithful chromosome segregation.

Here, we introduce a workflow based on a combination of quadruple-trap (Qtrap) optical tweezers, fluorescence microscopy, and microfluidics to manipulate the centromeric regions of isolated native human metaphase chromosomes. Using this assay, we determined the mechanical properties of centromeres before and after cohesin cleavage, over a broad range of forces, and find that cohesin is an important determinant of centromere stiffness at high forces, but its cleavage does not trigger chromosome segregation *in vitro*. To capture these mechanical characteristics of centromeres, we developed a “softened Hierarchical Worm-Like Chain” (sHWLC) model. Finally, we observed that after cohesin cleavage, the mitotic DNA decatenase, Topoisomerase IIα (TopIIα), is selectively recruited to centromeres under pulling forces up to 400 pN. Centromeric DNA entanglements are thus likely an additional cohesion factor that prevents segregation. Our experimental workflow paves the way to systematically investigate the mechanics and regulation of chromosome segregation.

## Chromosome attachment and handling

To enable direct mechanical measurements on centromeres using our Qtrap optical tweezers workflow, we specifically biotinylated the outer kinetochores and telomeres of human chromosomes. We first fused biotin ligase BirA* to the outer kinetochore protein HEC1 and to Telomeric Repeat-binding Factor 1 (TRF1) in modified HCT116 cells (**Fig. S1** and **Methods**)(*32*, *35*). BirA* biotinylates proteins in its proximity, allowing attachment of optically trapped streptavidin-coated microspheres to individual chromosomes at the centromeres and telomeres(*32*, *36*). Cells were first synchronized by arresting them in G2 using the CDK1as system (see **Methods**). After release from G2, and in the presence of biotin, cells were subsequently arrested in metaphase by inhibiting the Anaphase Promoting Complex/Cyclosome (APC/C) using proTAME-Apcin(*37*). Metaphase chromosomes were then isolated and purified as described previously(*32*, *36*) (**Fig. 1a** and **Methods**).

**Fig. 1.**
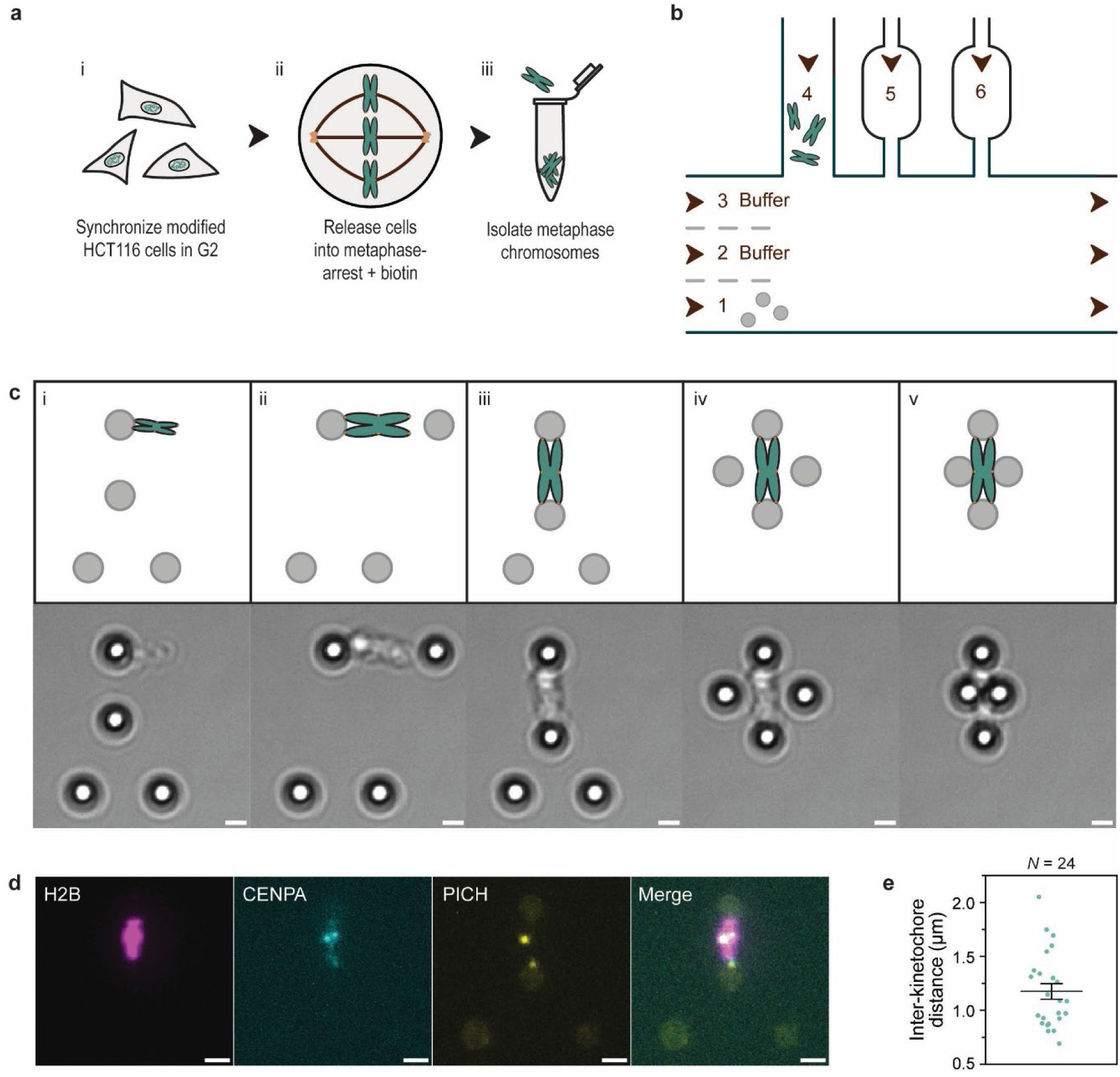
Centromeric attachment of metaphase chromosomes using quadruple-trap optical tweezers. **a,** Cell culture and chromosome isolation. Cells were first synchronized in G2 (16 h), after which they were released into metaphase arrest (4 h) in the presence of biotin. Metaphase chromosomes were then isolated and purified further (see **Methods**). **b,** Overview of microfluidic flow-cell. Four beads were trapped in channel 1 and subsequently steered to channel 4, which contains chromosomes. After catching an individual chromosome with one of the beads, the beads and chromosome were moved to channel 3 to attach the remaining beads to the other telomeric end and the centromeres. Properly oriented chromosomes could either stay in channel 3 or be moved to a side channel (channel 5 or 6) with a large reservoir and narrow entry, allowing long incubation in different buffers of interest. **c,** Attachment of beads to a trapped chromosome. After trapping a chromosome at one telomeric end in channel 4 (i), the beads and chromosome were moved to channel 3. A flow was then applied to the chromosome to stretch it laterally. The second bead was then attached to the other telomeric end (ii) and the chromosome was brought to a vertical orientation (iii). This allowed the other two beads to approach (iv) and attach to the outer kinetochores (v). **d,** Verification of centromere location by fluorescent markers. Fluorescence images were taken prior to attachment of the second bead pair to prevent fluorophore bleaching caused by the nearby trapping lasers. Scale bars indicate 2 µm. **e,** Inter-kinetochore distances of chromosomes attached by four beads, measured at a centromeric tension of 50 pN. Error bars represent SEM.

Chromosome trapping and handling was carried out within a six-channel laminar flow-cell integrated with the optical tweezers, allowing swift buffer exchange and extended incubations in large reservoir-shaped side channels (**Fig. 1b**). To trap a chromosome, four optically trapped beads were moved to side channel 4 containing biotinylated chromosomes. There, a chromosome was attached to the uppermost bead by one of its telomeric ends (**Fig. 1c** and **Movie S1**). The beads and chromosome were then moved back to buffer channel 3, where a gentle flow was applied to stretch the chromosome sideways, allowing attachment of the second bead to the other telomere. This telomeric attachment was necessary to stretch the chromosome, exposing its centromeric regions, and allowing attachment of the other bead pair to the outer kinetochores. The location of the centromere regions was usually apparent from chromosome rotation during flow-stretching (**Movie S2**) and brightfield images prior to attachment of the second bead pair (**Fig. 1c**). Additionally, the location of the centromeres could be verified by fluorescence imaging of the centromeric histone variant mCherry-CENP-A and mNeon-PICH, a protein recruited to DNA bridges that occur at centromeres (**Fig. 1d**)(*17*). While holding the chromosome taut with the telomere-attached beads, the remaining two beads were gently pushed into the identified centromeric regions to attach them there (**Fig. 1c**). After proper attachment of the chromosome, the force on the telomeric bead pair was minimized for subsequent experiments. In this Q-trapped configuration, the force applied to the centromeres could be controlled and monitored with piconewton precision. The observed mean inter-kinetochore distance of 1.2 ± 0.1 µm, measured at a centromeric tension of 50 pN, is consistent with previously reported values of bioriented human chromosomes (**Fig. 1e**)(*38*, *39*).

### Mechanics of metaphase centromeres

To characterize the mechanical properties of metaphase centromeres, we first trapped and attached chromosomes using Qtrap optical tweezers as described above. We then applied increasing force to the centromeres up to 250 pN by moving one bead attached to the centromere away from the other with a constant velocity of 0.1 µm/s (**Fig. 2a,b** and **Movie S3**), mimicking the force loading geometry of the mitotic spindle *in vivo*. At these forces, the extension of centromeres was reversible (**Fig. S2a**), as was shown previously for telomeric stretching of chromosomes using optical tweezers and micropipette aspiration(*30–32*). Although centromeres showed heterogeneous stretching behaviour, dramatic stiffening at high forces was evident in all measurements (**Fig. 2b**).

**Fig. 2.**
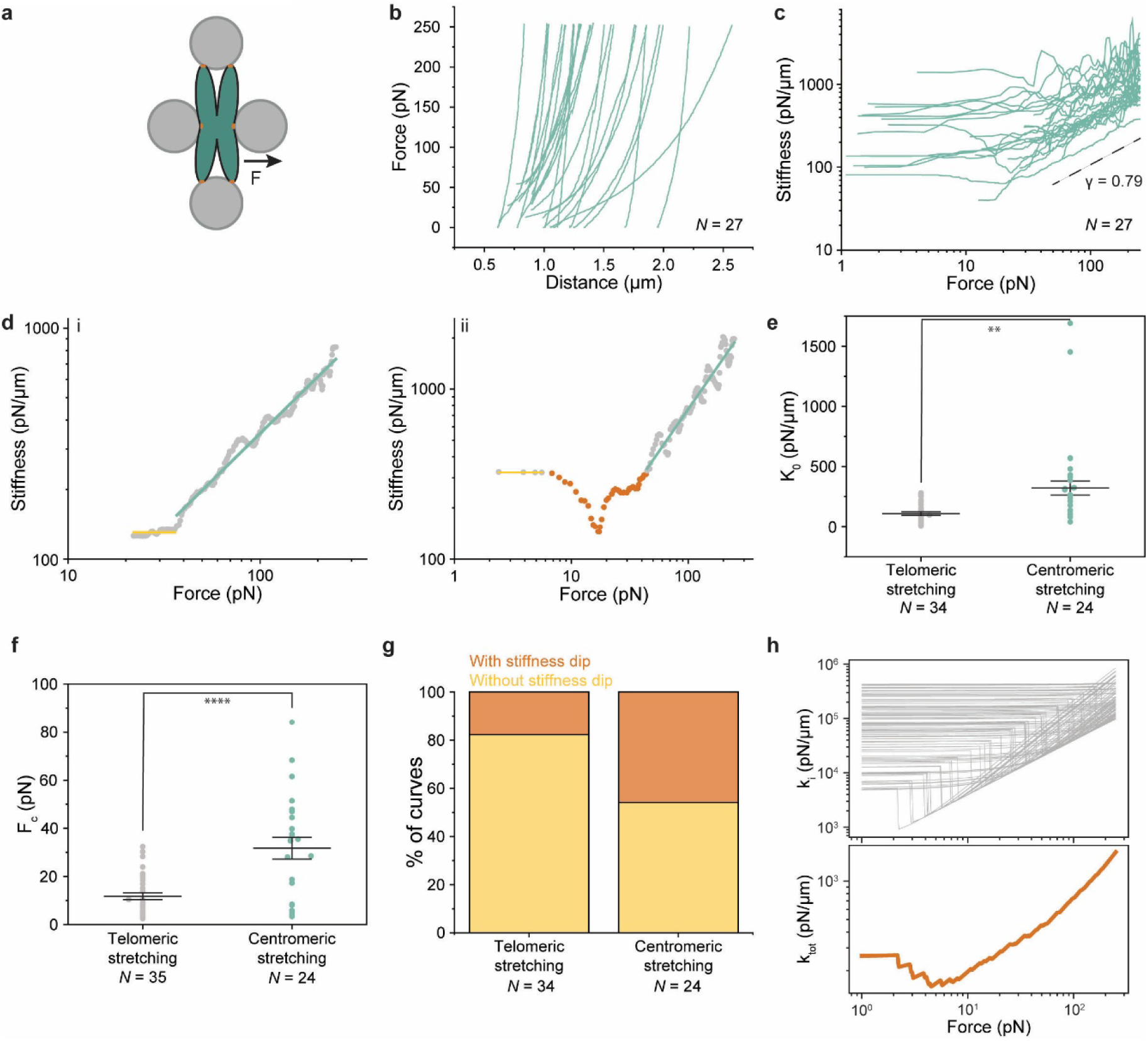
Mechanical characterization of metaphase centromeres. **a,** A metaphase chromosome held by four beads attached to its telomeres and centromeres. A stretching force is applied to the centromeric beads by moving the rightmost centromeric bead to the right with a constant velocity. **b,** Force-distance curves collected during the stretching of centromeres. **c,** Stiffness curves derived from panel b (blue), and their mean stiffening exponent *γ* (dashed black). **d,** Examples of analysis of stiffness curves (i) consisting of an initial linear stiffness (yellow), followed by a regime where the stiffness follows power law scaling (blue), or (ii) where a dip in stiffness (orange) is found between the linear and power-law regimes. **e,** Initial stiffness *K*_0_ of whole chromosomes stretched by their telomeres(*32*) (grey) and centromeric stretching (blue). **f,** Critical force *F_c_* of whole chromosomes stretched by their telomeres(*32*) (grey) and centromeric stretching (blue). **g,** Quantification of occurrence of stiffness dips for telomeric stretching of whole chromosomes and centromeric stretching **h, s**HWLC model. Upper panel: tangent stiffness *K_i_* of individual WLC elements. Lower panel: total assembly stiffness *K_tot_* (orange). ** *p* < 0.01, **** *p* < 0.0001. *N* represents number of chromosomes. Error bars represent SEM.

To quantify the stiffening behaviour of centromeres, we determined the stiffness *K* by calculating the derivative of the force with respect to the distance and used it to generate stiffness-force plots (**Fig. 2c**). The centromeres exhibited a linear initial stiffness *K*_0_ at low forces, and a stiffening regime at high forces beyond the critical force *F_c_*, where the stiffness follows a power-law dependence, *K* ∼ *F^γγ^* (**Fig. 2d**), as was shown for longitudinally stretched chromosomes with beads attached to the telomeres(*32*). To quantitatively analyse the mechanical properties of centromeres and compare them with the behaviour of longitudinally stretched chromosomes, we extracted *K*_0_, *F_c_*, and stiffening exponent *γ* from both centromeric (this study) and telomeric (Meijering *et al*.(*32*)) stretching experiments (see **Methods**). Centromeres showed a significantly higher *K*_0_ than longitudinally stretched chromosomes (**Fig. 2e** and **Table S1**). This difference can be explained by size differences between whole chromosomes and centromeres, as we found that the Young’s moduli of centromeres and whole chromosomes are similar (**Fig. S3a** and **Table S2**, see **Methods**), in agreement with a recent AFM study(*40*). The critical force *F_c_*, on the other hand, was significantly higher for centromeric stretching compared to telomeric stretching (**Fig. 2f** and **Table S1**), indicating that centromeres commence stiffening at higher forces than whole chromosomes. The three times higher *K*_0_ and *F_c_* might be important for centromeres to prevent irreversible damage in response to pulling forces from the mitotic spindle.

Above the *F_c_*, centromere stiffness follows a power-law scaling, with an average stiffening exponent *γ* of 0.79 ± 0.04 (**Fig. 2c** and **Table S1**), which deviates from classical polymer models such as the freely jointed chain (FJC; *γ* = 2) or the worm-like-chain model (WLC; *γ* = 1.5)(*41*, *42*), but is nearly identical to the stiffening exponent calculated from whole chromosome stretching (0.82 ± 0.05)(*32*). Such anomalous stiffening behaviour can be explained by the hierarchical worm-like-chain (HWLC) model(*32*), where *γ* < 1 is the result of sequential stiffening of heterogenous components of the chromosome, each of which acting as a WLC element with individual stiffening properties. The similarity in stiffening exponents between centromeres and whole chromosomes suggests that the stiffening response of centromeres can be attributed to sequential stiffening of underlying centromeric structures.

Interestingly, a fraction of centromeres exhibited a pronounced dip in stiffness between the linear and power-law scaling regimes (**Fig. 2d(ii)**). This nonlinear softening response was reversible (**Fig. S2b**) and not due to the presence of the four traps around the chromosome or minor force fluctuations caused by overlapping interference patterns of beads in close proximity (**Fig. S2c,d**)(*43*). Such stiffness dips were also present in a fraction of the telomeric chromosome stretching data from Meijering *et al*.(*32*), albeit with a less frequent occurrence (**Fig. 2g**) but with similar dip width and depth compared to centromeric stretching (**Fig. S3b-d**). Together, this indicates that a dip in stiffness between the linear and power-law scaling regime is a feature of some chromosomes, regardless of the stretching direction.

The presence of stiffness dips cannot be explained by the original HWLC model(*32*). To account for the dips in stiffness in our theoretical model, we developed a softened HWLC (sHWLC) model, which includes a single modification for all underlying WLC elements: at a set threshold force (0.2 *F_c_*_,*i*_), a proportion of the elements (*p*_soft_; 0.5) decrease in stiffness by an order of magnitude (see **Methods**). Such a decrease in element stiffness could be interpreted as a force-induced reversible conformational change occurring in a chromosomal component. With this minimal modification, the sHWLC model effectively recapitulates the emergent dips in stiffness occurring at intermediate forces (**Fig. 2h**).

### Cohesin cleavage alters centromere mechanics

To test the impact of cohesin on centromere mechanics, we set out to cleave the cohesin present on isolated metaphase chromosomes *in vitro*. To this end, we introduced three consecutive TEV protease cleavage sites (3xTEV) into an unstructured region of the cohesin subunit RAD21/Scc1 (see **Methods**), an approach that has previously been used to trigger chromosome segregation in *Drosophila*(*44*). Transfection of TEV protease into metaphase-arrested HCT116 RAD21-3xTEV cells induced chromosome segregation within 15 min (**Fig. S4a**). However, when TEV protease was added to isolated metaphase chromosomes from the same cell line, chromosome segregation did not occur (**Fig. S4b**), despite near-complete cleavage of cohesin under these conditions (**Fig. S4c-d**).

We further tested if pulling forces are required to segregate sister chromatids after cohesin cleavage. We first preincubated chromosomes in a buffer ± TEV protease in bulk for 30 min and introduced them into the Qtrap optical tweezers instrument. We then stretched their centromeres up to 250 pN with a constant pulling velocity (0.1 µm/s). TEV protease mediated cohesin cleavage induced no notable difference in chromosome morphology or inter-kinetochore distances under pulling forces (**Fig. 3a-c**, **Movie S4,** and **Fig. S5**). The overall stiffening response of centromeres after TEV protease mediated cohesin cleavage was also similar to that of native metaphase centromeres (**Figs. 3d** and **2c**). Additionally, there were no statistically significant changes in the initial stiffness *K*_0_ or Young’s modulus *E* (**Fig. 3e**, **Fig. S3a**, and **Table S1**). However, both the critical force *F_c_* and stiffening exponent *γ* increased significantly after cohesin cleavage (**Fig. 3f,g** and **Table S1**). The increased stiffening exponent suggests a change in the distribution of force-bearing elements, according to the HWLC model(*32*). In addition, we observed an increase in the prevalence of stiffness dips (**Fig. 3h**). Therefore, cohesin is a major contributor to the mechanical integrity of centromeres at high forces.

**Fig. 3.**
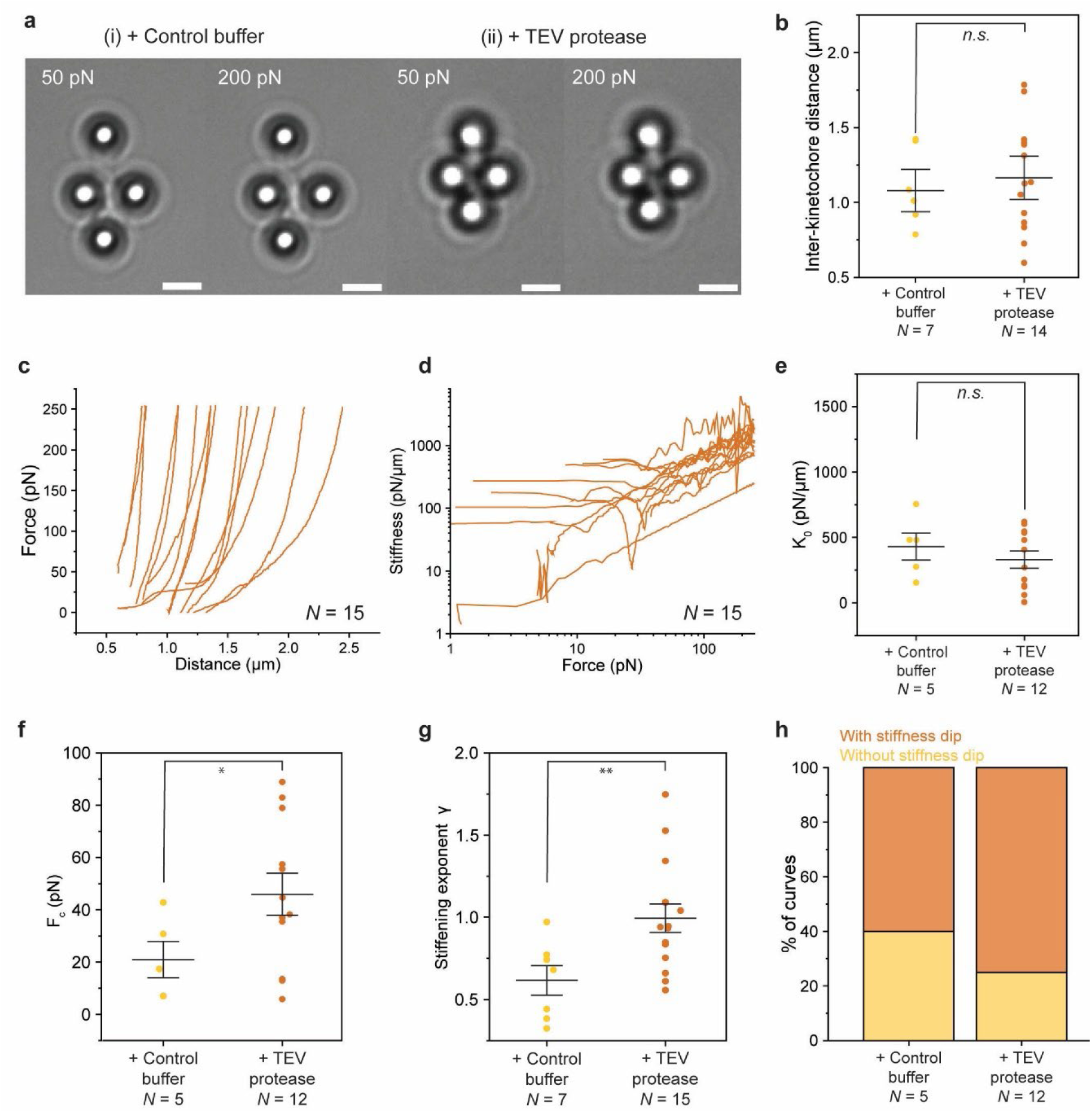
Mechanical changes in the centromere following cohesin cleavage. **a,** Brightfield images of optically trapped metaphase chromosomes ± TEV-protease pretreatment. Scale bars indicate 2 µm. **b,** Inter-kinetochore distances (1.08 ± 0.09 *vs* 1.2 ± 0.1 µm), measured at a centromeric tension of 50 pN, of metaphase chromosomes that were incubated with control buffer (yellow), or incubated with TEV protease (orange) prior to optical tweezers experiments, respectively. **c,** Stretch curves of centromeres of individual chromosomes following TEV-protease incubation. **d,** Stiffness curves of individual chromosomes from panel c (orange). **e-g,** Initial stiffness *K*_0_ (panel e), critical force *F_c_* (panel f) and stiffening exponent *γ* (panel g) of centromeres of individual metaphase chromosomes that were incubated with control buffer (yellow) or incubated with TEV protease (orange), respectively. **h,** Quantification of occurrence of stiffness dips. *n.s*. not significant, * *p* < 0.05, ** *p* < 0.01. *N* represents the number of chromosomes. Error bars represent SEM.

The sHWLC model introduced above not only recapitulates stiffness dips, it also predicts an increase in critical force *F_c_* and stiffening exponent *γ* compared to the HWLC model (**Fig. S6a**). This indeed matches the observed differences in centromere mechanics following cohesin cleavage (**Fig. 3f,g** and **Table S1**). To explain the difference in the frequency of stiffness dips following cohesin cleavage, we optimized the sHWLC model by fine-tuning the proportion of elements that undergo softening, *p*_soft_ (**Fig. S6b,c**). At a lower *p*_soft_, stiffness dips occurred only occasionally, when by chance elements with a low initial stiffness softened. Thus, an increase in prevalence of stiffness dips could be explained in the sHWLC model as an increase in *p*_soft_. We find that cohesin cleavage leads to a two-fold increase in the proportion of underlying centromeric elements that can undergo softening (**Fig. S6c,** see **Methods**). Despite the complexity of centromere structure, the relatively simple sHWLC model faithfully captures the mechanical properties of centromeres before and after cohesin cleavage.

### TopIIα binds centromeres under tension

Although cohesin cleavage changes the mechanical properties of centromeres, it does not lead to chromosome segregation *in vitro*, which implies that other centromeric components still hold the sister chromatids together and prevent chromosome segregation. The most likely candidate for such a component is remaining DNA entanglements, which are known to play a role in sister chromatid cohesion(*13*, *14*, *16*, *21*). After TEV protease mediated cohesin cleavage, we incubated Q-trapped chromosomes in a buffer reservoir containing the DNA intercalating dye YO-PRO-1 and indeed observed a substantial YO-PRO-1 signal at the centromeres upon application of stretching forces (6/6 chromosomes; **Fig. 4a** and **Movie S5**). This suggests that there were still many short DNA entanglements present that need to be resolved before chromosome segregation can occur.

**Fig. 4.**
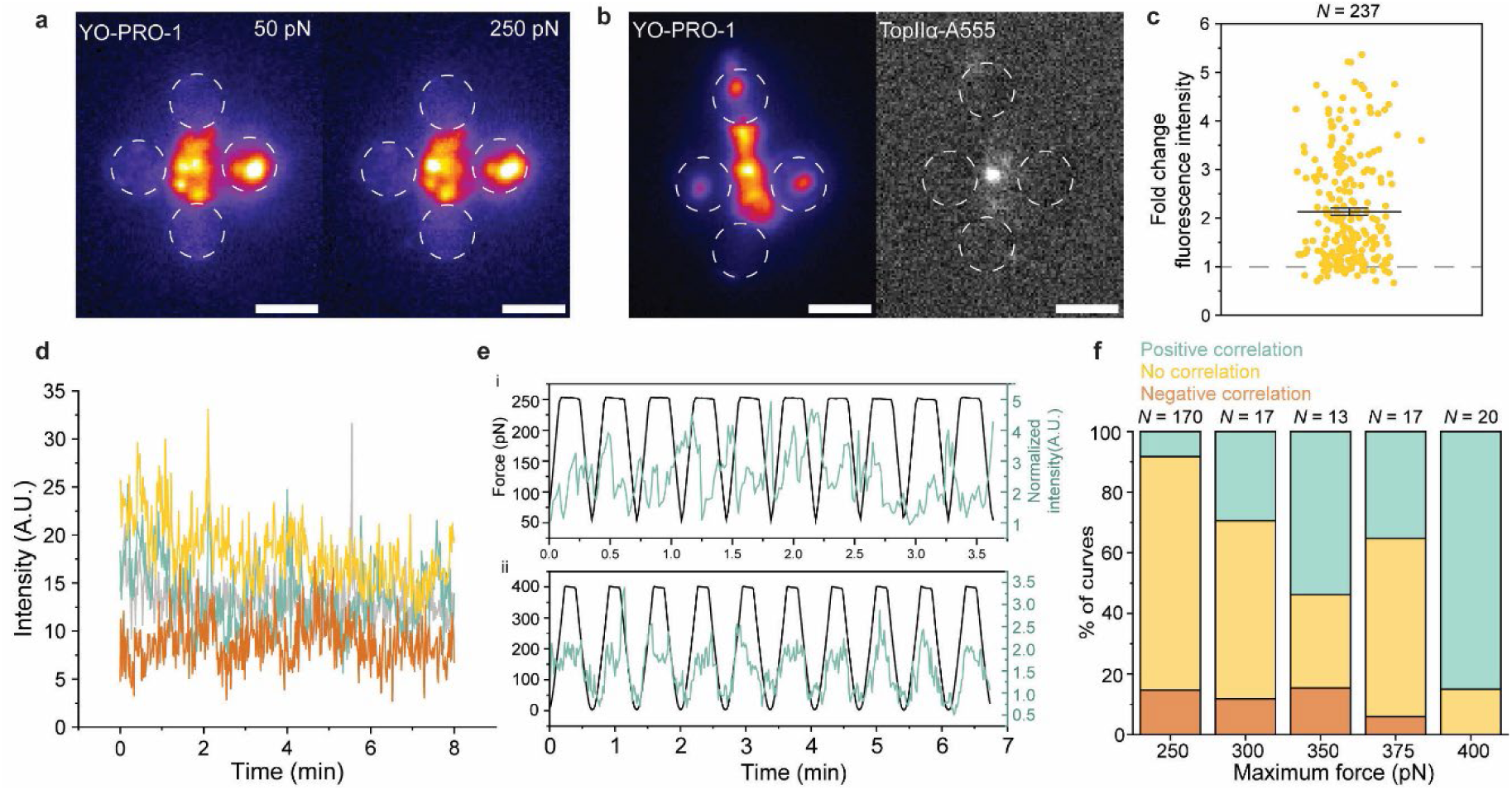
Topoisomerase IIα localizes selectively to centromeres under tension after cohesin cleavage. **a,** Example of chromosome morphology during centromeric stretching in the presence of YO-PRO-1 intercalator at 50 pN (left) and 250 pN (right). **b,** Example of YO-PRO-1 signal from a chromosome prior to TopIIα incubation (left) and the same chromosome later, showing concentration of TopIIα-A555 signal at the centromere. Both images were taken at a centromeric tension of 250 pN. Scale bar indicates 2 µm. **c,** Fold change of TopIIα-A555 fluorescence signal of centromeres relative to chromosome arms shows that centromeres have a higher average fluorescence signal per pixel than chromosome arms in 211/237 stretch cycles from four chromosomes. Dotted line indicates normalized fluorescence signal from chromosome arms. *N* represents the number of stretch cycles. **d,** Centromere TopIIα-A555 signal during 400 pN force clamp. Many TopIIα-A555 molecules stay bound, and peaks indicate binding of new molecules. Colours indicate different (*N* = 4) chromosomes. **e,** Example of 10 stretch cycles up to 250 pN (i), showing limited correlation, and up to 400 pN (ii), showing positive correlation of TopIIα-A555 signal to the applied centromeric tension. **f,** Quantification of Pearson’s correlation coefficient of fluorescence intensity with maximum applied force. *N* represents the number of stretch cycles. Scale bars indicate 2 µm. Error bars represent SEM.

The major factor involved in removal of centromeric DNA entanglements during mitosis is TopIIα, an enzyme that is present on centromeric DNA intertwines into early anaphase and is necessary for their timely decatenation(*13*, *45*). The reported low force tolerance of TopIIα(*46–48*) seemingly conflicts with its role in mitosis, where spindle forces are thought to be omnipresent. Thus, we investigated the influence of centromeric tension on TopIIα binding. To this end, we confirmed the activity of TopIIα fluorescently labelled with Alexa555 (A555) using a bulk DNA decatenation assay (**Fig. S7a**, see **Methods**). Isolated metaphase chromosomes were then incubated in bulk with TEV protease, trapped with Qtrap optical tweezers as described before, and stretched in a side channel containing YO-PRO-1. The trapped chromosomes were subsequently moved to another side channel containing TopIIα-A555 to monitor the binding behaviour of TopIIα under centromeric tension using fluorescence microscopy. TopIIα preferentially bound to the (peri)centromere region compared to the chromosome arms (**Fig. 4b,c**, **Fig. S7b**, and **Movie S6**). At a constant centromeric pulling force of 400 pN, the A555 fluorescence intensity at the centromeres overall remained stable over the course of the measurement (8 min), and new fluorescence peaks appeared (**Fig. 4d**). Hence, despite the high force applied to the centromeres, TopIIα binding persists.

Next, multiple stretch cycles were performed sequentially with a maximum centromeric force ranging from 250 to 400 pN, after which the Pearson’s correlation coefficient between the applied force and centromeric fluorescence signal was determined for each stretch cycle (see **Methods**). This analysis shows that at a maximum force of 250 pN, most cycles (77% of *N* = 170) show no significant correlation between force and fluorescence intensity (**Fig. 4e(i),f**). This indicates that binding of TopIIα to centromeres is not affected by centromeric forces up to 250 pN. Unexpectedly, when the maximum stretch force was increased to 400 pN, the percentage of curves showing a positive correlation increased (84% of *N* = 20; **Fig. 4e(ii),f**), suggesting that TopIIα binding to centromeres is stimulated by increasing centromeric tension up to 400 pN. Binding of TopIIα to centromeres, however, did not lead to significant chromosome morphology changes, centromere lengthening, or chromosome segregation (**Fig. S7c,d**). Thus, the force-stimulated TopIIα binding is likely an essential intermediate in the complete decatenation of centromeric DNA entanglements.

## Discussion

Here, we present a quadruple-trap optical tweezers assay to study the mechanical properties of native human mitotic centromeres. The stiffening behaviour of metaphase centromeres overall resembles whole chromosomes(*32*). In particular, the stiffening exponent *γ* of centromeres is nearly identical to longitudinally stretched chromosomes, implying a similar chromatin organization despite a perpendicular pulling direction. Chromosome arms are proposed to consist of bottle-brush structures(*49*). Therefore, our results indicate that centromeres also consist of bottle-brush structures, oriented perpendicularly to chromosome arms, in support of recent reports(*29*, *50–52*).

Cohesin cleavage does not lead to chromosome segregation *in vitro*, but it is a major contributor to centromere mechanics, as its cleavage increases the critical force *F_c_*, stiffening exponent *γ*, and the prevalence of stiffness dips. To gain mechanistic insights into the origin of stiffness dips, we developed a model (sHWLC) by adding a softening transition to the original HWLC model(*32*). Remarkably, the introduction of this softening transition also correctly predicts the increased critical force *F_c_* and stiffening exponent *γ* after cohesin cleavage. Furthermore, the increase in prevalence of stiffness dips after cohesin cleavage can be explained by a higher *p*_soft_, the proportion of underlying elements that undergo softening. The structural basis of this softening transition is still unclear. We hypothesise that mechanical instabilities, such as buckling, near centromeres could give rise to the observed dips in centromere stiffness (**Supplementary text** and **Fig. S8**).

Remaining centromeric DNA entanglements seem to prevent chromosome segregation following cohesin cleavage in our assay. Notably, we observed accumulation of TopIIα at (peri)centromeric regions at high centromeric forces (250-400 pN), which is in the range of the reported force generating capacity of the mitotic spindle(*23*, *24*, *26*). As there are multiple parallel DNA catenanes at the centromere, the pulling force will be distributed. Therefore, the force on each DNA catenane is likely within the active force range of TopIIα(*46–48*). Still, this TopIIα binding did not lead to chromosome segregation. Efficient resolution of centromeric DNA entanglements may therefore require additional factors. One candidate of such a factor is condensin, which is highly enriched at centromeres(*29*, *53*) and guides TopIIα towards decatenation(*54*, *55*). Nevertheless, the high force binding of TopIIα is probably essential for effective chromosome segregation. Finally, we anticipate that our single-molecule approach presented here can be expanded upon to disentangle the complex process of chromosome segregation as well as many other mechanisms relevant to centromeres and cell division in health and disease.

## Acknowledgments

We thank Andreas Biebricher for help with DNA intercalators and Janni Harju for help with data interpretation.

## Author contributions

Conceptualization: IDH, GJLW

Methodology: TVMC, CFN, BH

Investigation: TVMC, CFN, ZL, BH, RC, CCC

Visualization: TVMC, ZL

Funding acquisition: IDH, GJLW

Supervision: CB, IDH, GJLW

Writing – original draft: TVMC, ZL, GJLW Writing – review & editing: all authors

## Data, code, and materials availability

The data supporting the findings of this study will be available at a data repository prior to publication. The code used to analyse centromere stiffness and to construct the modified HWLC model and buckling model will be available from a code repository prior to publication

## Supplementary Materials

## Materials and Methods

### Cell line modifications

All transfections were done using the Neon electroporation system (Thermo Fisher Scientific), according to the manufacturer’s protocol. All oligos used in this study are listed in **Table S2**.

### Synchronisation and mitotic arrest

We employed a chemical genetics approach to create HCT116 cells that could be efficiently arrested in G2 and released into mitosis. In short, endogenous CDK1 was knocked out by CRISPR and the ATP analogue sensitive CDK1 mutant CDK1as knocked in, as described previously(*32*, *56*, *57*). Plasmids for CDK1as engineering were kind gifts from William Earnshaw (Addgene 118596 and 118597)(*56*) and Zsuzsanna Izsvak (Addgene 34879)(*58*).

### Enabling in vitro cohesin cleavage

To enable cohesin cleavage by addition of TEV protease, we inserted 3xTEV sites into the unstructured region of endogenous RAD21 (exon 7). For this, an ALT-R HDR donor oligo with the 3xTEV sequence flanked by homology arms mutated to not be recognised by the gRNA was synthesised (IDT). A gRNA crRNA targeting exon 7 of RAD21 was designed using the Custom Alt-R^™^ CRISPR-Cas9 guide RNA tool and synthesised (IDT). The crRNA was annealed with tracrRNA-ATTO550 and combined with ALT-R Cas9 enzyme (IDT) to make RNP complexes. RNP complexes and ALT-R HDR donor oligos were transfected into HCT116 CDK1as H2B-emiRFP-670 cells and transfected cells were treated with V2 HDR enhancer (IDT) for 18 hours. Transfectants were single cell sorted for ATTO550 signal using a Wolf Cell sorter (Nanocellect). Correct clones were detected by genomic PCR and confirmed by immunoblotting.

### Fluorescent fusion proteins

To fluorescently tag H2B with emiRFP-670, the HCT116 CDK1as cells were transfected with pH2B-emiRFP670 (Addgene 136571), a kind gift from Vladislav Verkhusha(*59*). Clones were selected by treatment with 1.5 mg/mL G418 (Sigma-Aldrich) and visual inspection.

Endogenous PICH (gene name ERCC6L) was fluorescently tagged at its N-terminal with mNEON. To create a homology directed repair vector we cloned a ERCC6L 5’ homology arm, mNEON gene without stop codon, linker, then ERCC6L 3’ homology arm into the pUC19 backbone vector (Addgene 50005). The 266bp 5’ homology DNA oligo was amplified by genomic PCR and a DNA oligo containing the mNEON gene flanked by a short linker sequence and a 197 bp 3’ homology arm was synthesised (IDT). A short sequence (CAAGAGTC) was added in before the mNEON gene to recreate a mutated 5’ UTR not recognised by the gRNA. These two DNA fragments were designed to have a 25 nt overlap with adjoining fragments and were assembled into the pUC19 vector digested with EcoRI by HiFi NEBuilder DNA assembly (NEB). A gRNA crRNA targeting the N terminal of the ERCC6L gene was designed, synthesized, and assembled into RNP complexes as described above. ERCC6L gRNA RNP complexes and pUC19-mNEON-ERCC6L HDR vector were transfected into HCT116 CDK1as H2B-emiRFP-670 RAD21-3xTEV cells. Transfected cells were treated with V2 HDR enhancer (IDT) for 18 hours. Clones were selected by single cell flow cytometry sorting for ATTO550 and mNEON using a Wolf Cell sorter (Nanocellect) and confirmed by visual inspection and genomic PCR.

Ectopic expression of mCherry-CENP-A was added to the HCT116 CDK1as RAD21-3xTEV H2B-emiRFP-670 mNEON-PICH cells by transfection with the pLenti6-CENP-A-mCherry plasmid, which was a gift from Torsten Wittmann (Addgene 89767)(*60*). Positive clones were selected by treatment with 7.5 µg/mL Blasticidin (Sigma-Aldrich) and confirmed by visual inspection.

### Biotinylation of telomeres and centromeres

Production of lentivirus for biotinylation of telomeres by expression of BirA-TRF1 was performed as described previously(*32*). For biotinylation of centromeres by expression of BirA*-HEC1, the TRF1 gene was excised from the transfer plasmid by digestion with NheI and KpnI restriction enzymes (NEB), and the HEC1 gene inserted by HiFI NEBuilder DNA assembly (NEB). Here, BirA* refers to a mutant of BirA that is used for proximity labelling. Successful biotinylation of centromeres was confirmed by immunofluorescence (See below; **Fig. S1**). For biotinylation of both telomeres and centromeres, cells were transduced with equal dose of lentivirus containing either transfer plasmid BirA-TRF1 or BirA-HEC1.

### Cell culture and chromosome isolation

HCT116 RAD21-3xTEV CDK1as H2B-iRFP mNEON-PICH mCherry-CENP-A BirA*-TRF1 BirA*-HEC1 cells were grown in high-glucose DMEM (Gibco) supplemented with 10% fetal bovine serum (FBS; Gibco) and penicillin-streptomycin (Gibco), in a humidified incubator set at 37°C, 5% CO_2_. Cells were synchronized by blocking in G2 for 16 hours with 0.25 µM 1NM-PP1 (Sigma-Aldich). Subsequently, they were released into mitosis for four hours with 12.5 µM proTAME (Cayman Chemicals) and 25 µM Apcin (Sigma-Aldrich) to stall them in metaphase(*37*, *45*), and 12.2 mg l^-1^ biotin (Sigma-Aldrich) to biotinylate centromeres and telomeres. For YO-PRO-1 and TopIIα experiments (see below), mitotic arrest was extended to six hours, in an effort to reduce cohesion between sister chromatids(*37*, *45*).

Chromosomes were isolated from metaphase-stalled cells using a polyamine buffer method(*61*), as described previously(*32*, *36*). In short, metaphase cells were harvested via gentle mitotic flush-off and subjected to a swelling buffer consisting of 5 mM Tris-HCl (pH 8.0) and 75 mM KCl. Cells were washed and resuspended in isolation buffer (15 mM Tris-HCl (pH 8.0), 2 mM EDTA, 0.5 mM EGTA, 80 mM KCl, 20 mM NaCl, 0.5 mM spermidine, 0.2 mM spermine and 0.2% Tween-20), supplemented with cOmplete Mini Protease and PhosSTOP Phosphatase Inhibitor Cocktails (Roche). Cell lysis was achieved by 25 strokes of a Dounce homogenizer with a tight-fitting pestle. After clearing the lysate of cell debris, chromosomes were purified by centrifugation over a glycerol step-gradient containing two layers (30% and 60% (v/v) glycerol, in isolation buffer). Chromosomes were collected from the 60% glycerol fraction and stored at -20°C. Isolated chromosomes could be used for at least several months without showing signs of degradation.

### Bulk experiments

#### Immunofluorescence and microscopy

Metaphase cells were harvested as described above. To assess TEV protease mediated cohesin cleavage in live cells, this cell suspension was then transfected ± TEV protease (NEB) at a concentration of 0.6 u/µL according to the manufacturer’s protocols using the Neon Electroporation system. Cells were then seeded and fixed as described below. To assess biotinylation of centromeres, harvested metaphase cells were added directly to 37°C media on Poly-L-lysine (Sigma-Aldrich) coated glass coverslips and seeded for a minimum of 15 min before fixation. Fixation and immunofluorescent staining of cells was performed according to the co-extraction method described by Bizard *et al.*(*62*). In brief, cells were washed with PBS then simultaneously fixed and permeabilized using PTEMF buffer (4% PFA, 0.25% triton-x 100, 20 mM PIPES pH 6.8, 1 mM MgCl_2_, 10 mM EGTA; Sigma-Aldrich). Hereafter washes and antibody staining was performed as previously described(*62*). Antibodies used were CENP-B (Santa Cruz, sc-32285, 1:200), PICH (Cell Signalling, 8886S, 1:100) and CREST (Antibodies Inc, 15-234, 1:500). Biotin was stained using Streptavidin-Alexa Fluor 568 (Invitrogen, 10297422) DNA was stained with DAPI (Sigma-Aldrich).

To assess the effect of TEV protease mediated cohesin cleavage on native chromosomes, isolated chromosomes were incubated ± 1 u/µL TEV protease (NEB) for 30 min at 30°C in polyamine buffer (20 mM HEPES pH 7.4, 0.5 mM EDTA, 80 mM KCl, 1 mM DTT, 1 mM spermidine, 0.4 mM spermine, 0.1% (v/v) Tween-20). The treated chromosomes were seeded onto poly-L-Lysin coated coverslips for 15 min on ice. The coverslips were then washed twice in polyamine buffer without Tween-20, stained with DAPI, and mounted on microscopy slides in polyamine buffer in Vectashield mounting medium without DAPI (H-1200, Vector Laboratories). Slides were kept at 4°C and imaged within three days of mounting.

Microscopy was performed using a Zewass confocal laser scanning microscope, ZEWASS LSM 900, equipped with an Airyscan2 detection unit. The NA oil immersion alpha Plan-Apochromat 63X/1.40 Oil was used, with Zen Blue software 3.5. Comparative images of the blue, green and red channels were captured using the Airyscan super-resolution mode. Images were taken in 12 bits, with a sequential, unidirectional acquisition, averaged on two images, with an image size of 1024 × 1024 pixels (pixel size 20 nm), a zoom of 3, pinhole 1 AU for confocal. with 405, 488 and 568 nm lasers at an intensity of 1%.

#### Immunoblotting

To confirm complete TEV protease mediated cohesin cleavage, 200 µL of isolated chromosome solution was diluted 3x in polyamine buffer (see above) ± 0.2 or 1 u/µL TEV protease (NEB) and incubated at 30°C or 27°C, for 30 min or 5 min, respectively. The chromosomes were then centrifuged at 3000 RCF for 15 min and resuspended in RIPA buffer with cOmplete Protease Inhibitor cocktail (Roche). Samples were incubated 10 min on ice before being sonicated for ten 30 sec on/off cycles in a water cooled sonicator at 4°C (Diagenode). Samples were then centrifuged at 21000 RCF for 15 min, after which the supernatant was moved to precooled tubes. These samples were used for standard immunoblotting as described previously(*63*), using the following antibodies: RAD21 (Abcam, ab-154769, 1:1000) Histone H3 (Abcam, ab1791, 1:1000)

#### Topoisomerase IIα purification and testing

Expression and purification of TopIIα was performed based on previously published methods(*64*). Briefly, a plasmid containing full-length human TopIIα (hTOP2a 12URAB), a kind gift from James Berger(*64*), was introduced into the *S. cerevisiae* strain INVsc1 (Invitrogen), using the LiAc/SS carrier DNA/PEG method(*65*). Starter cultures were incubated overnight at 30°C, 200 RPM in Yeast Synthetic Drop-out medium lacking uracil (Sigma), supplemented with 2% (w/v) glucose. Starter cultures were then diluted 1:20 in Yeast Synthetic Drop-out medium lacking uracil (Sigma), supplemented with 2% (v/v) sodium-D-L-lactate and 1.5% (v/v) glycerol and incubated overnight at 30°C, 200 RPM. At that point, the intermediate cultures were diluted 1:10 in Yeast Peptone medium supplemented with 2% (v/v) sodium-D-L-lactate and 1.5% (v/v) glycerol and incubated at 30°C, 200 RPM until an OD600 of 0.8-1.1 was reached. TopIIα expression was then induced by addition of 2% (w/v) galactose and incubation for an additional 6 hours at 30°C, 200 RPM. Cell pellets were harvested by centrifugation and stored at -20°C until use.

For purification, cell pellets were thawed on ice and resuspended in buffer A (20 mM Tris pH 8.5 at 4°C, 500 mM KCl. 20 mM Imidazole, 10% (v/v) glycerol, 1 mM DTT) supplemented with cOmplete Mini EDTA-free Protease Inhibitor tablets (1/10 mL, Roche). 5 mL of resuspension buffer was used per gram of cell pellet. Cells were lysed by 15-20 10 sec rounds of sonication, followed by 1 min cooling on ice. Lysate was cleared by centrifugation at 17000 RCF for 30 min and filtered using a 0.2 µm filter before being loaded onto a 5 mL Histrap HP column (Cytiva). The column was first washed with 40 column volumes of buffer A, and then with 10 column volumes of buffer B (20 mM Tris pH 8.5 at 4°C, 150 mM KCl. 20 mM Imidazole, 10% (v/v) glycerol, 1 mM DTT). Protein was eluted over a gradient of buffer C (20 mM Tris pH 8.5 at 4°C, 150 mM KCl. 200 mM Imidazole, 10% (v/v) glycerol, 1 mM DTT) and collected in 4 mL fractions. Fractions were assessed by SDS-PAGE. Then, fractions containing TopIIα were pooled, concentrated, and exchanged to storage buffer (20 mM Tris pH 8.5 at 4°C, 150 mM KCl, 20% (v/v) glycerol, 1 mM DTT) using Amicon Ultra centrifugal filters (30 KDa MWCO). Aliquots of concentrated protein were snap-frozen in liquid nitrogen and stored at -70°C.

To later visualize TopIIα with fluorescence microscopy, purified TopIIα was labelled with Alexa-555. To achieve this, purified protein was first dialyzed against labelling buffer (50 mM HEPES pH 7.0, 500 mM KCl, 10% (v/v) glycerol) overnight at 4°C. Then, purified protein was mixed at a 1:1 molar ratio with AlexaFluor-555 NHS Ester (Invitrogen) and incubated at 4°C for 10 hours. Afterwards, unbound dye was removed by buffer exchanging using Zeba Spin desalting columns (7K MWCO, Thermo Scientific) into storage buffer (see above). Aliquots of fluorescently labelled protein were snap-frozen in liquid nitrogen and stored at -70°C. With this method, a labelling ratio of 1.1 was achieved, indicating that on average, every TopIIα protein was labelled with 1.1 molecules of Alexa-555-NHS Ester.

Decatenation activity was tested using the Topogen Topoisomerase II Assay kit, according to the manufacturer’s protocol (**Fig. S7a**). 10 ng/µL kinetoplast DNA (kDNA) was incubated ± 30 nM TopIIα-A555 in a buffer consisting of 20 mM HEPES pH 7.4, 80 mM KCl, 1 mM spermidine, 0.4 mM spermine, 0.02% Tween-20, 1 mM DTT, 5 mM MgCl_2_, 1 mM ATP. Samples were incubated at 27°C for 5, 15 or 30 minutes.

### Quadruple-trap optical tweezers experiments

Trapping experiments were performed using a commercially available instrument that combines quadruple-trap optical tweezers with widefield fluorescence microscopy (C-trap, LUMICKS). The instrument was operated using Bluelake software (LUMICKS). Fluorescent imaging was achieved with a 638 nm excitation laser at 3%, a 561 nm excitation laser at 10%, and a 488 nm excitation laser at 10% (100% for visualizing YO-PRO-1), respectively. Excitation lasers were alternated to prevent bleed through to other detection channels.

Experiments were carried out within a six-channel flow-cell (**Fig. 1b**; Micronit) controlled by a microfluidic flow-system (LUMICKS). Prior to the onset of measurements, the flow-cell and microfluidic system were flushed with bleach, followed by sodium thiosulfate neutralization. To prevent chromosomes and proteins from sticking to the tubing and flow-cell surface, the system was passivated by overnight incubation with 0.1% casein in PBS, followed by rinsing with polyamine buffer (see above).

Prior to each measurement, the trap stiffness was calibrated using the thermal noise method and was usually around 2 pN/nm for the centromere-attached bead pair. Due to instrument limitations combined with the small bead size (see below), calibration of the telomere-attached bead pair was less stable, leading to an uncertainty in the force detected from this bead pair. Thus, no quantitative analysis was performed based on forces from this bead pair. After chromosome attachment, tension on telomeric beads was minimized during all force measurements to limit its effect on centromere mechanics. The temperature of the objective during measurements was around 27°C.

All channels contained polyamine buffer (see above). Additionally, channel 1 contained 1.25 µm or 1.76 µm streptavidin-coated polystyrene beads (Spherotech), and channel 4 contained 20-40 µL chromosome stock diluted in 500 µL polyamine buffer. For experiments where TEV-protease cleavage was done in bulk, outside of the optical tweezers, 20-40 µL chromosome stock was incubated in polyamine buffer supplemented with 1 mM DTT and ± 0.2 u/µL TEV protease (NEB) for 30 min at 30°C and then added to channel 4.

Bead trapping and positioning was automated using Bluelake software (LUMICKS). After trapping beads in channel 1, a chromosome was attached to the first bead by a telomeric end in channel 4. The trapped chromosome was then moved to channel 3 where it was flow-stretched to attach the other telomeric end to another bead. Then, while holding the chromosome taut, the remaining two beads were attached to the outer kinetochore. After attaching, tension on the telomeric bead pair was reduced to minimize its effect on centromere mechanics. Then, the centromeric region of the chromosome was stretched three times up to 250 pN and back by moving the right centromeric bead to the right. Chromosomes where the inter-telomere length was shorter than the inter-kinetochore distance were discarded, as they were likely attached improperly. Chromosomes where fluorescence signal from H2B-iRFP, mCherry-CENP-A and mNeon-PICH were absent were also discarded.

Then, for experiments where the chromosome was moved to a side channel for incubation, a tension of at least 50 pN was applied to the centromeric region prior to moving. This tension was required to prevent additional links between the beads and centromeric regions from forming, which could alter the stretching behaviour. For YO-PRO-1 experiments, channel 5 contained polyamine buffer supplemented with 5 nM YO-PRO-1. Chromosomes were moved to this channel and stretched repeatedly. For TopIIα stretching experiments, trapped chromosomes were first stretched in channel 5 containing YO-PRO-1. Then they were moved to channel 6, which contained ± 3 nM TopIIα in a buffer consisting of 20 mM HEPES pH 7.4, 80 mM KCl, 1 mM spermidine, 0.4 mM spermine, 0.02% (v/v) Tween-20, 1 mM DTT, 5 mM MgCl_2_, 1 mM ATP. Chromosomes were repeatedly stretched here, first to a maximum force of 250 pN, and sequentially up to a maximum force of 400 pN. Four chromosomes were stretched to a maximum force of 250 pN. Two of these were then further stretched up to 300, 350, 375 and 400 pN. After TopIIα incubation a final round of stretching was performed in channel 5. For force clamp experiments in the presence of TopIIα, trapped chromosomes were first stretched in channel 5 containing YO-PRO-1. Then they were moved to channel 6 containing TopIIα as before, and a constant force of 400 pN was directly applied to the centromeres for 10 min. The first two minutes of fluorescence data from each force clamp measurement was excluded from analysis as extensive TopIIα binding occurred upon channel entry.

### Data analysis

#### Centromere stiffness

Force data was exported using the Python package Pylake and analysed using OriginPro 2024b and custom-written Python code. Stiffness analysis was carried out on the third consecutive stretch of each chromosome, in a force range of 1 -250 pN. Force-distance curves were first smoothed using the locally weighted scatterplot smoothing (LOWESS) method(*66*), with a window size of 0.1. The differential stiffness *K* was then calculated by numerical differentiation of smoothed force with respect to distance and subsequently plotted against force *F* on a log-log scale. Data points showing drastic jumps due to presence of ruptures in force-distance curves, were excluded from further analysis.

In custom written python code, the stiffness curves could then be classified into different regimes (**Fig. 2d**). If present, a stiffness plateau at low force (< 50 pN), was first determined, initiated by manual selection of a plateau datapoint. The average value of this plateau signifies the initial linear stiffness *K*_0_. After the linear regime, a dip in stiffness sometimes followed. A stiffness dip was assigned when a stiffness value was lower than *K*_0_ minus three standard deviations. The end of a dip was marked by the start of the power law scaling regime. This power law scaling regime followed the stiffness dip if present, or otherwise the linear regime. This regime started when a stiffness value larger than *K*_0_ plus three standard deviations was found. The stiffening regime was fitted using *K* = *aF^γγ^*, where *a* is the scaling factor and *γ* is the stiffening exponent. For some stiffness curves, mostly those that started at a slightly higher force, only the power law scaling regime could be detected. This regime was then fitted as described above, from the first data point.

To consider the effect of geometrical properties of chromosomes/centromeres on the *K*_0_, we calculated the Young’s Modulus 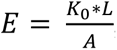, where the length *L* is defined as the chromosome length for whole chromosomes and the inter-kinetochore distance for centromeres (both at 100 pN), and the area *A* was set to 1 µm^2^ for both.

#### Image analysis

Image analysis was done using ImageJ. Fluorescence images of H2B-iRFP, mCherry-CENP-A and mNeon-PICH were only taken before attaching the centromeric bead pair. This was because after attaching the centromeric beads, the fluorophores could bleach quickly due to the close proximity of the trapping lasers, thus deforming their signal.

All fluorescence quantification was done on background corrected signal. To quantify TopIIα-A555 fluorescence, the chromosome was first segmented into upper arm, lower arm, and centromere regions based on CENP-A and PICH location before bead attachment, combined with YO-PRO-1 and brightfield images (**Fig. S7b**). To determine the fold change of TopIIα-A555 fluorescence signal in centromeres *vs* chromosome arms, the total fluorescence intensity per pixel in the centromere region was divided by the total fluorescence intensity per pixel in chromosome arms, for each stretch cycle of each chromosome.

To correlate the observed centromeric TopIIα-A555 fluorescence signal with the applied force, the force data was first binned to 1 Hz to match the framerate of the fluorescence imaging. Then, the

Pearson’s correlation coefficient (PCC) between the centromeric fluorescence signal and applied force was determined for each stretch cycle of each chromosome. The resulting correlations were then scored as significantly positive (PCC > 0, *p* < 0.05), neutral (*p* ≥ 0.05), or significantly negative (PCC < 0, *p* < 0.05).

#### Statistics

Statistical significance of differences in mean values between two groups were determined by two-sample T-test (two-sided) with equal variances not assumed (Welch correction). For analysis between three or more groups one-way ANOVA with Tukey’s HSD post-hoc testing was used.

### sHWLC model

Following Meijering *et al.*(*32*), the centromere was modeled as an assembly of *N_sub_* = 100 worm-like chain elements connected in series. Individual elements are modeled as inextensible worm-like chains (WLCs) using the Marko–Siggia force–extension relation

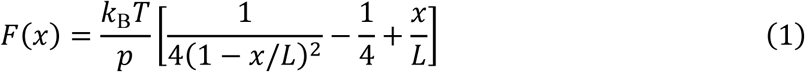

where x is the end-to-end extension, *L* the contour length, *p* the persistence length, and k*_B_*T = 4.114 pN · nm. For a given applied force, the extension of each element was obtained numerically, and the total extension was computed as

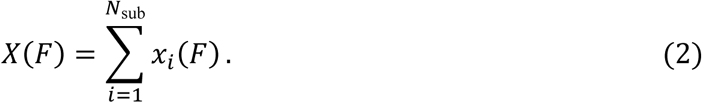

The effective stiffness of the assembly was defined as the tangent stiffness

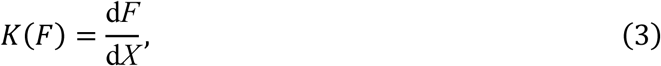

evaluated numerically from the force–extension curve. Each element was assigned a characteristic force scale *f_c,i_* drawn independently from a truncated power-law distribution 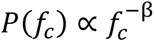 with *β* = 0.79 and bounds 10–1000 pN. The linear-response stiffness was related to the force scale through

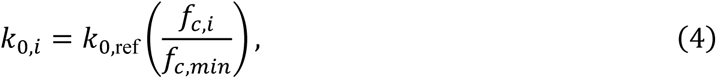

with *α* = 1 and *k*_0,ref_ = 300 pN/μm. The persistence length of each WLC was chosen such that the small-extension limit of the WLC matched this prescribed linear stiffness,

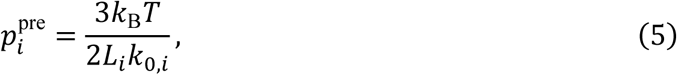

where all elements had equal contour length *L_i_* = *L*_tot_/*N*_sub_ with *L*_tot_ = ∼1 μm. At high forces, the HWLC framework exhibits emergent power-law stiffening, with exponent *γ* = *α* + *β* − 1.

To model the cohesin-cleaved condition, elements were allowed to undergo a reversible force-induced softening transition. Before stretching, a fraction of the elements was given the ability to transition to a softer state above a threshold force

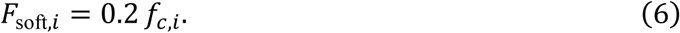

Which elements were able to soften was determined randomly for each individual element with a probability *p*_soft_ = 0.5. Softening was implemented by increasing the persistence length by a factor of ten,

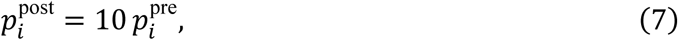

which reduces the tangent stiffness of the WLC. The post-transition force–extension relation was shifted to ensure continuity of X(*F*) at *F*_soft,*i*_. Force–extension curves were evaluated on a logarithmically spaced force grid spanning 1–250 pN. High-force stiffening was quantified by fitting the asymptotic regime of *K*(*F*) to a power law *K*(*F*) ∼ *AF*^γ^.

To quantify how often the sHWLC model produces an observable stiffness dip, the simulation was repeated for many independently initialized chromosome pulls at fixed values of *p*_soft_. Here, *p*_soft_ should be interpreted as the average proportion of WLC elements that are able to soften for a given simulated chromosome pull. For each realization, the subset of softened elements was initialized once at the start of the pull and then kept fixed throughout that realization, such that no stochasticity was present during pulling.

For the dip frequency analysis, repeated realizations were performed over a range of *p*_soft_ values, with *N_sub_* = 50 and *k*_0,ref_ = 450 pN/μm. For each realization, the total effective stiffness was computed. Curves were then classified based on the appearance of a stiffness dip, which was defined as a > 20% reduction in stiffness compared to *k*_0_. This was used to determine the dip frequency at each *p*_soft_.

### Centromere buckling model

To explore geometry-driven softening, we constructed a two-dimensional spring network whose topology approximates the experimentally observed centromere geometry (**Fig. S8a**). Nodes represent effective attachment sites along chromatin, and connections represent loop-mediated interactions between segments. Black springs form the main chromosome arms, yellow springs denote DNA entanglements, and blue springs represent additional cohesin-mediated connections that brace the centromeric region. Each connection has rest length *L*_0_ determined by the initial geometry and current length *L*. For compression (*L* ≤ *L*_0_), the connection behaves as a Hookean spring,

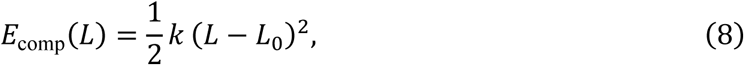

representing an effective compressive resistance arising from steric and crosslink constraints. In extension, connections exhibit nonlinear stiffening inspired by the Marko–Siggia worm-like chain form. Defining the reduced extension

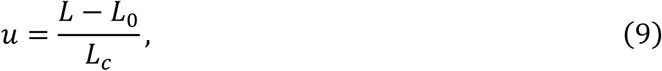

Where *L_c_* is an available contour-extension scale proportional to *L*_0_, the extension energy is

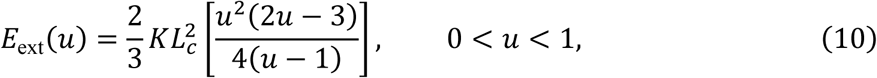

with configurations satisfying *u* ≥ 1 forbidden. This choice ensures linear response at small extension and strong stiffening as contour length is approached.

The network is constrained to mimic the experimental pulling geometry: nodes at the top, bottom, and left are fixed, while nodes on the right are displaced. For each displacement step, all unconstrained nodes were relaxed by minimizing the total potential energy

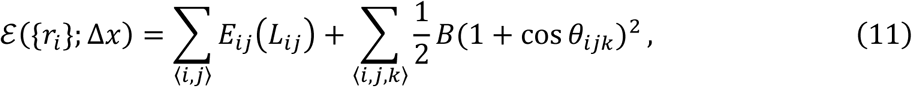

where {*r_i_*} is the collection of node coordinates, *E_iii_* denotes the energy of each connection and the second term sums over all node centred triplets formed by pairs of springs sharing a node. This penalises hinge-like folding, as to prevent buckling from occurring at near-zero force. Equilibrium corresponds to vanishing net forces on all unconstrained nodes, obtained numerically by energy minimization, where the previous state configuration was used as initial condition.

For the fixed-end geometry, top, bottom, and left-side anchor nodes were held fixed while right-side centromeric nodes were displaced. For the free-end geometry, the top and bottom chromosome-arm endpoints were released while keeping the centromeric pull geometry unchanged. Spring stiffnesses were assigned by spring type, with higher stiffness in the bridges representing many parallel connections. Effective stiffness is extracted from the resulting force– extension curve using tangent stiffness. Simulations used displacement-controlled quasistatic pulling over 0.4 µm. Reaction forces were then computed from the minimized configurations. An overview of the simulation parameters used can be found in **Table S3**.

## Supplementary Text

### Centromeric buckling model

Although the modified HWLC model presented in this study recapitulates multiple features of observed centromere stiffness, it does not suggest what might cause the sudden softening of underlying elements. Elastic instabilities, such as buckling, are one of the few possible phenomena within a polymer network that could explain reversible softening in response to pulling forces. Though it might seem contradictory, in a complex network under stretching forces geometric considerations imply that sub-components of the network could be under compressive forces and could thus buckle. Mechanically, buckling introduces an additional deformation pathway. Rather than increasing extension solely through stretching of individual elements, the structure can extend through geometric rearrangements caused by the buckling of compressed elements.

This mechanism can result in a stiffness dip: additional extension is achieved through conformational change rather than increased tension. At higher forces, the structure straightens and elements approach their nonlinear stiffening regime, leading to an increase in effective stiffness. This process is geometric, reversible, and force-dependent, making it a potential candidate for the element softening mechanism in our sHWLC model. We thus wondered if buckling was possible within centromeres, and if so, could cause the observed dips in stiffness following the low force regime.

To investigate this, we constructed a minimal model motivated by recent insights on centromere structure and organization(*29*, *67*); human centromeres have been shown to exhibit a bipartite organization where the centromere of each chromatid consists of two functional units, and the two subdomains are likely held together by cohesin(*29*), In our model, the chromosome is represented as a 2D WLC-spring network where nodes represent coarse-grained chromatin attachment sites and springs represent effective DNA loop-mediated connections with a set initial stiffness (**Fig. S8a**; see **Methods**). The stiffnesses of the entanglements were set to be larger as a way to coarse grain multiple parallel entanglements. Bipartite centromeres within each sister chromatid are connected by cohesin-mediated DNA entanglements, whereas the sister chromatids are connected by cohesin-mediated DNA entanglements as well as cohesin-independent DNA entanglements. In this network, the right-side nodes were displaced and the reaction force was computed from energy minimised configurations, matching our experimental setup. Then, the effective stiffness can be 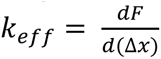

Running this simulation shows that buckling can occur both before and after cohesin cleavage, leading to pronounced softening in both cases (**Fig. S8b**). *In vivo,* a pulling force is applied to both centromeres and telomeres are untethered. Thus, we next mimicked the *in vivo* network configuration and again applied a pulling force to the network (**Fig. S8c**). In this configuration, buckling and the resulting stiffness dips still occurred both before and after cohesin cleavage. Thus, by building a minimal 2D model mimicking proposed centromere geometry, we show that buckling within chromosomes is plausible in both *in vitro* and *in vivo* scenarios and could explain the dips in stiffness observed in centromere stretching experiments.

Conceptually, this network model complements the sHWLC description. In the sHWLC model, the stiffness dip arises from reversible softening of individual elements within a heterogeneous hierarchy. Here, an analogous macroscopic response emerges from collective geometric reorganization. Both pictures introduce a reversible, force-dependent pathway that transiently increases compliance before nonlinear stiffening dominates at higher forces.

**Fig. S1.**
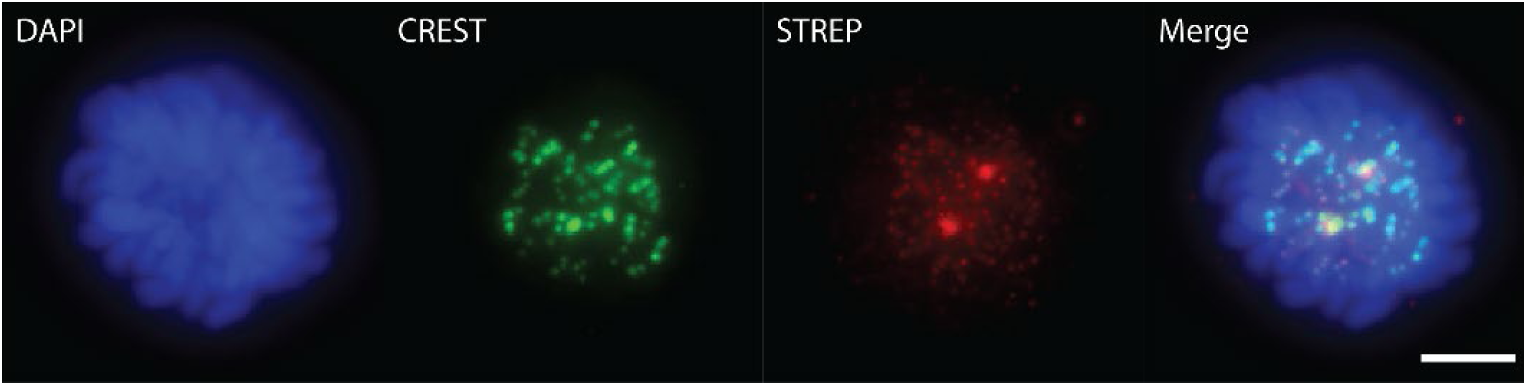
Biotin co-localizes with the centromere marker CREST in mitotic cells. DNA is stained with DAPI (blue). Centromere is visualized using anti-centromere antibody CREST (green), Biotin is visualized with streptavidin-Alexa Fluor 568 (red). Scale bar indicates 5 µm.

**Fig. S2.**
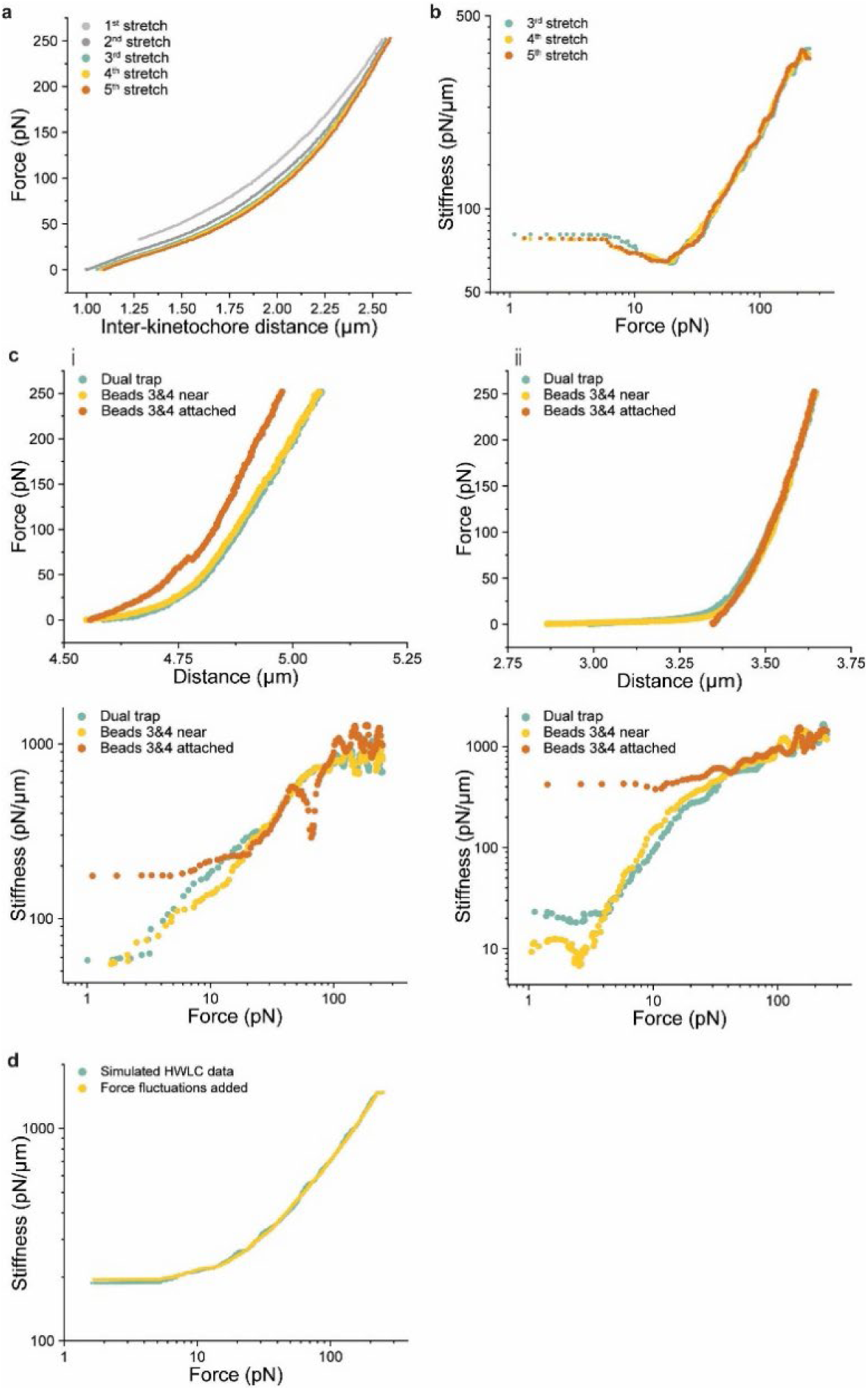
Stiffness dips are reversible properties of chromosomes. **a,** Example of five stretch cycles on the same metaphase chromosome. Stretching becomes fully reversible from the third stretch cycle onwards. **b,** Differential stiffnesses of stretch curves from a. This shows that dips in stiffness reoccur in subsequent stretch cycles and are thus reversible. **c,** Stretch (top) and corresponding stiffness curves (bottom) of chromosomes stretched by their telomeres only (blue), with a second bead pair near the centromeres (yellow), and after attaching the second bead pair to the centromeres (orange). An initial dip in stiffness is not caused by the proximity of the second bead pair. It is either absent (i) or present (ii) in the initial stretch and stays that way. *N* = 3 chromosomes. **d,** Example of simulated HWLC data (blue), overlayed with force fluctuations caused by small inter-bead distances (yellow), measured in absence of any sample between the beads. These force fluctuations did not cause stiffness dips in any of *N* = 15 stretch cycles from *N* = 5 bead pairs.

**Fig. S3.**
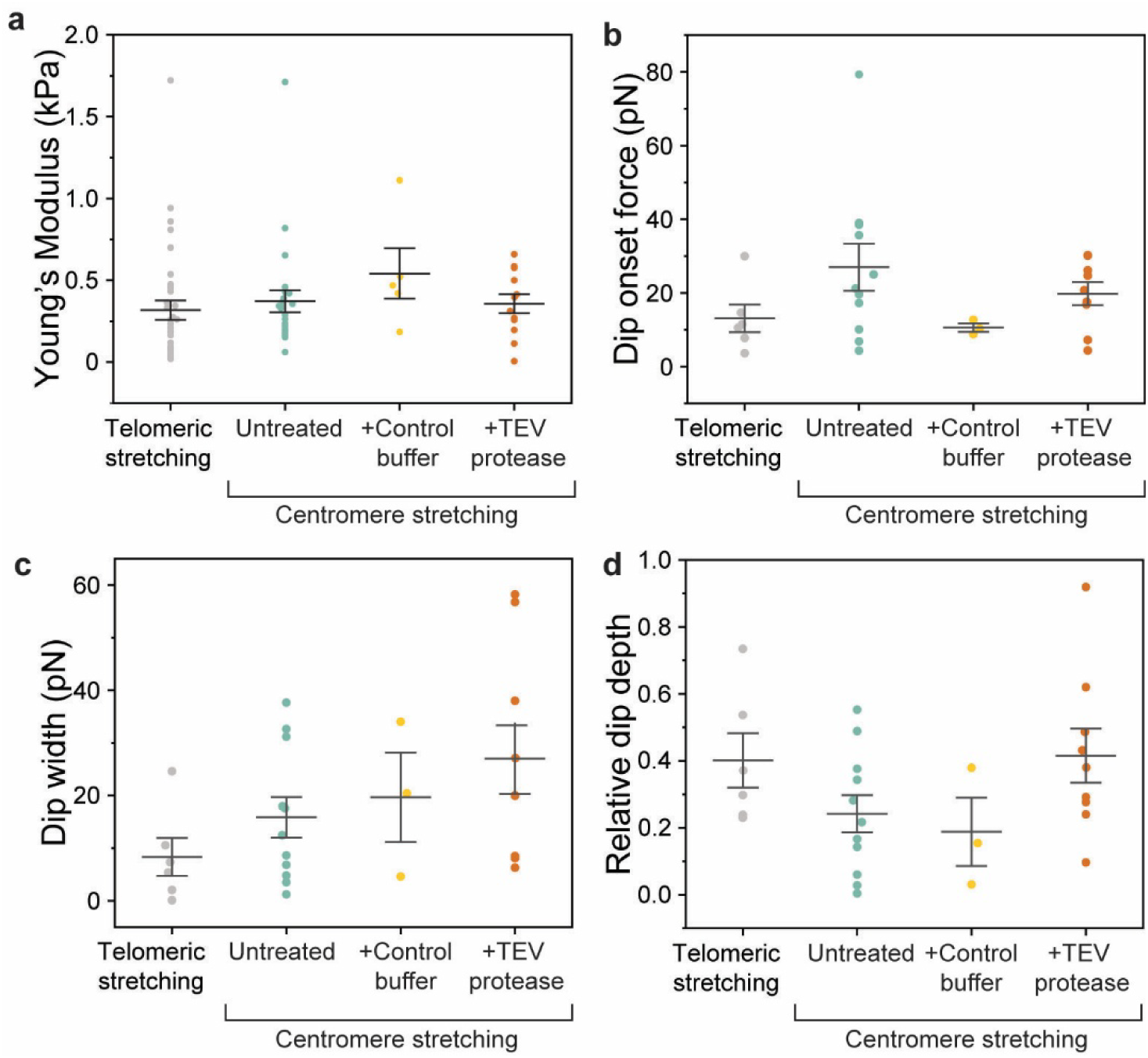
Additional centromere stiffness quantification. **a**, Young’s moduli. *N* = 35, 24, 5, 12 for telomeric stretching(*32*), and metaphase, control incubation and TEV protease incubation, respectively. **b-d:** Quantification of observed dips in stiffness following the linear stiffness regime. **b,** Dip onset force. **c,** Dip width. **d,** Depth of the stiffness dip relative to *K*_0_. *N* = 6, 11, 3, 9 for telomeric stretching(*32*), centromeric stretching of untreated metaphase chromosomes, control incubation, and TEV protease incubation, respectively. No statistically significant differences were found (one-way ANOVA). Error bars represent SEM.

**Fig. S4.**
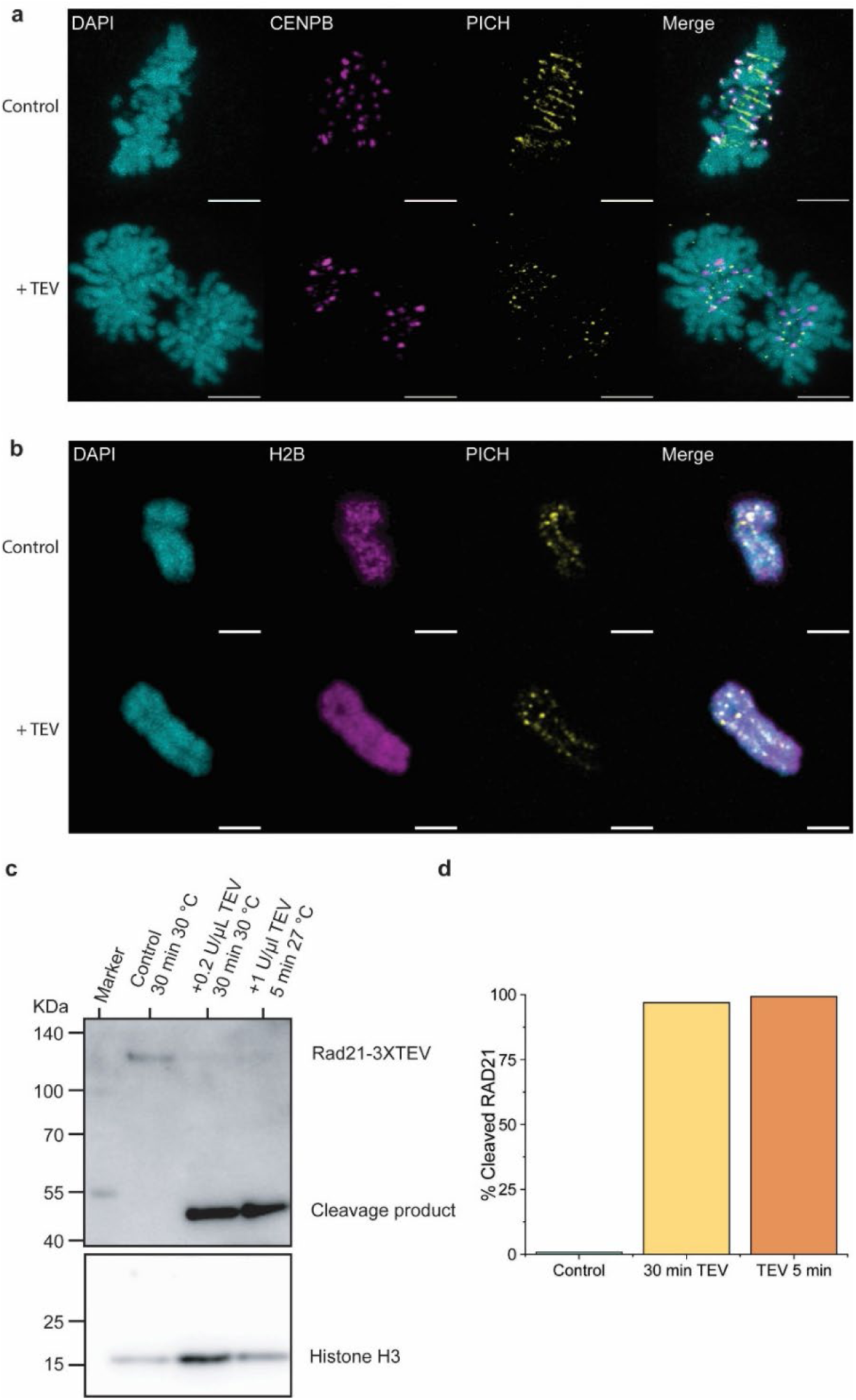
TEV protease-induced cohesin cleavage. **a,** Metaphase-arrested cells carrying 3xTEV cleavage sites in cohesin subunit SCC1 remain stalled in metaphase after electroporation with control buffer (top), but progress to anaphase after electroporation with TEV protease (bottom). *N* = 100 cells for each of three biological replicates for each condition. For those, segregation was observed for 4, 5, 7 cells in the absence of TEV protease, and 95, 96 and 98 cells incubated with TEV protease, respectively. Scale bar indicates 5 µm. **b,** In chromosomes isolated from metaphase-arrested cells, addition of TEV protease does not result in chromosome segregation. Sister chromatids remained connected in *N* = 16/16, 17/17 cases for control and TEV incubation, respectively. Scale bars indicate 2 µm. **c, d** Western blot showing TEV protease-induced cleavage of the cohesin subunit RAD21 (**c**) and its quantification (**d**).

**Fig. S5.**
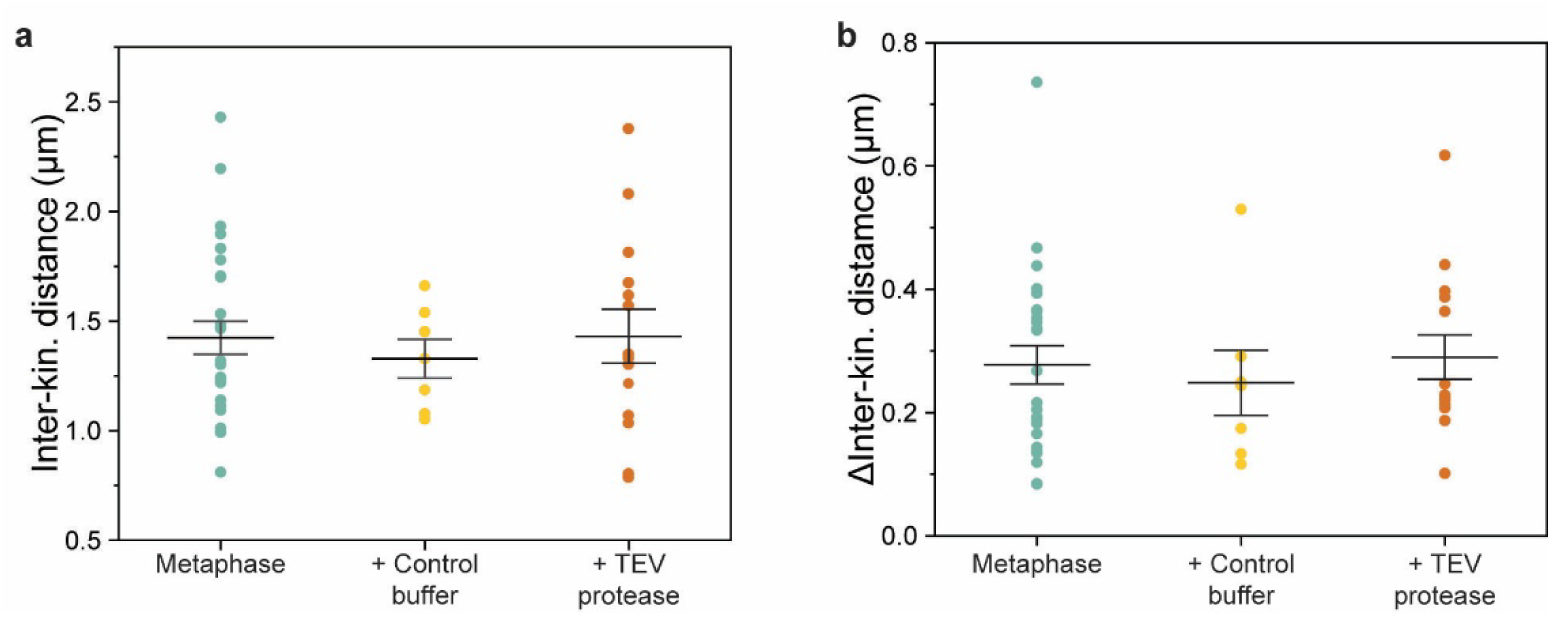
Inter-kinetochore distance does not change following cohesin cleavage. **a,** Inter-kinetochore distance at a centromeric force of 200 pN. N=27, 7, 14, respectively. **b,** ΔInter-kinetochore distance measured between centromeric forces of 50 and 200 pN. *N* = 24, 7, 14, respectively. No statistically significant differences were found (one-way ANOVA). Error bars represent SEM.

**Fig. S6.**
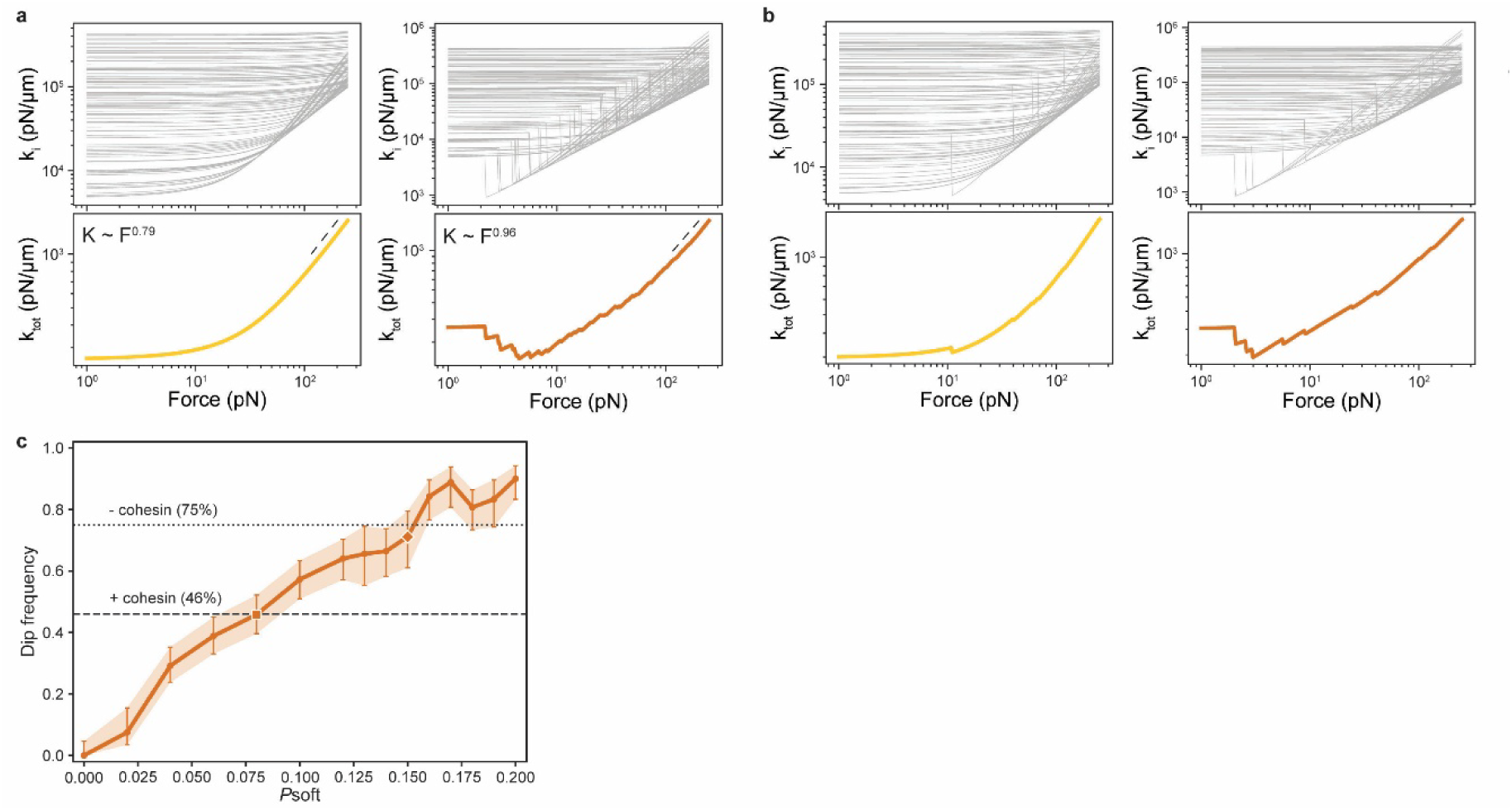
sHWLC model. **a,** Original HWLC model (i) and sHWLC model (ii). Upper panels: tangent stiffness *K_i_* of individual WLC elements. Lower panels: total assembly stiffness *K_tot_* (yellow/orange) and high-force fits of stiffening exponents (black dotted lines). **b,** At a *p*_soft_ of 0.1, in some iterations no visible stiffness dip appears (i). In other iterations, only if the elements that soften happen to have a low *F_c_*, a stiffness dip is clearly visible (ii). Upper panels: tangent stiffness *K_i_* of individual WLC elements. Lower panels: total assembly stiffness *K_tot_* (yellow/orange) **c,** Dip frequency in the modified HWLC model as a function of *p*_soft_. A dip frequency obtained from repeated realizations of the modified HWLC model as a function of *p*_soft_ for a dip definition of 20% reduction relative to the initial stiffness. Horizontal dashed lines indicate the experimentally observed dip fractions of 46% (+ cohesin) and 75% (-cohesin). Error bars indicate the 95% binomial finite-sample confidence interval.

**Fig. S7.**
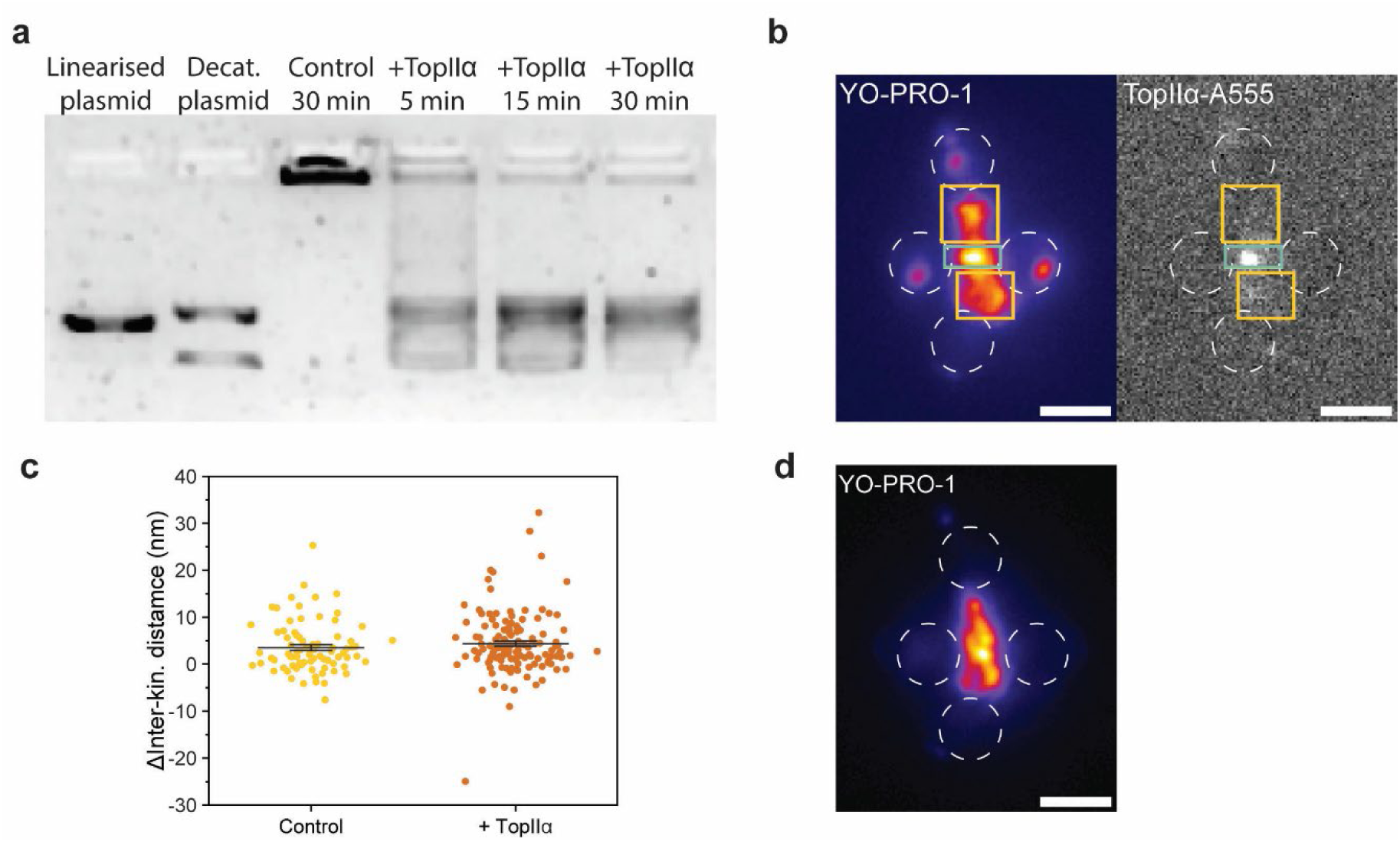
TopIIα experiments. **a,** TopIIα-A555 decatenation assay. kDNA plasmids that have been linearized (lane 1) and decatenated (lane 2). Catenated kDNA plasmid remains stuck in the well (lane 3; buffer control incubation). Incubation of catenated kDNA with TopIIα-A555 results in decatenation (lane 4-6). Smearing of DNA in lanes 4-6 is due to the presence of polyamines in the used buffer (see **Methods**). **b,** Example of ROIs used to define centromere (blue) and chromosome arms (yellow) for TopIIα fluorescence analysis. ROIs were found using YO-PRO-1 signal, combined with signal of mCherry-CENP-A and mNeon-PICH prior to attaching the centromeric beads, and visual inspection of brightfield images of chromosome spinning (see **Movie S2**). **c,** Quantification of lengthening during incubation with control buffer (yellow) or TopIIα (orange). Length was determined at a centromeric force of 200 pN. Mean lengthening per stretch cycle was 3.4 ± 0.6 vs 4.3 ± 0.6 nm, respectively. *N*= 80, 138 stretch cycles from *N* = 4, 7 chromosomes, respectively. **d,** YO-PRO-1 signal of chromosome (same as Fig. 4b) after TopIIα incubation. Image taken at centromeric tension of 80 pN. Scale bars indicate 2 µm. Error bars represent SEM.

**Fig. S8.**
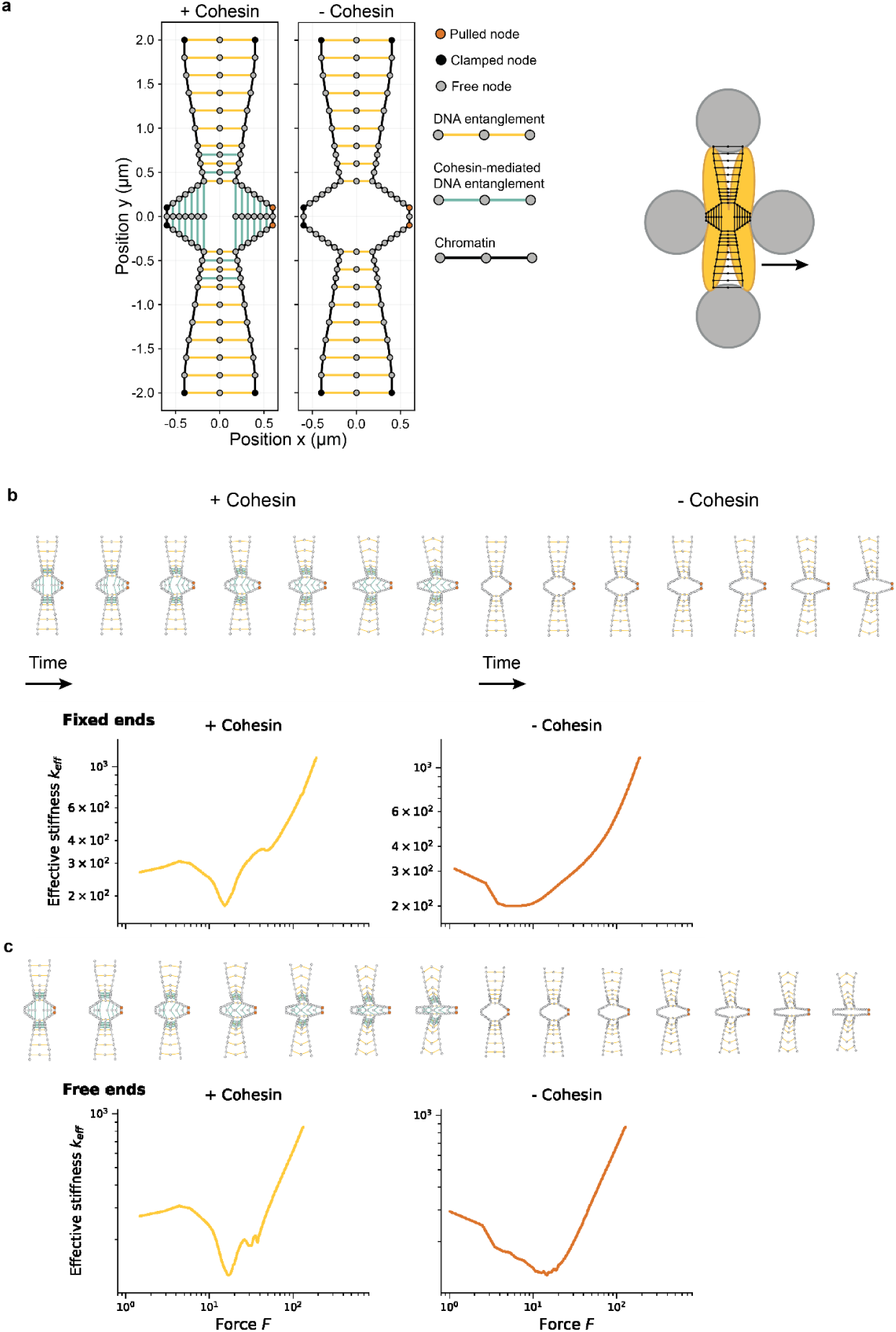
Buckling within centromeres could be the cause of observed dips in stiffness. **a,** Simplified representation of chromosome geometry as a 2D spring network. **b,** The effective stiffness *k_eff_* calculated from pulling on the network mimicking our experimental setup in metaphase (left) and after cohesin cleavage (right). Upper panels show change in network geometry over time. **c,** The effective stiffness *k_eff_* calculated from pulling on the network representing *in vivo* geometry, where telomeric ends are unconstrained, in metaphase (left) and after cohesin cleavage (right). Upper panels show change in network geometry over time.

**Table S1.** Mechanical parameters of chromosomes and centromeres.

| | | $K_0$<br>(pN/μm) | $F_c$ (pN) | $\gamma$ | $E$ (kPa) |
| --- | --- | --- | --- | --- | --- |
| Telomeric stretching(32) | | $110 \pm 20$ | $12 \pm 1$ | $0.82 \pm 0.05$ | $320 \pm 60$ |
| Centromeric stretching | Untreated | $320 \pm 60$ | $32 \pm 5$ | $0.79 \pm 0.04$ | $370 \pm 70$ |
| | Incubation with control buffer | $430 \pm 100$ | $21 \pm 7$ | $0.6 \pm 0.1$ | $540 \pm 150$ |
| | Cohesin cleavage by TEV-protease incubation | $330 \pm 70$ | $46 \pm 8$ | $1.0 \pm 0.1$ | $360 \pm 60$ |

**Table S2.**
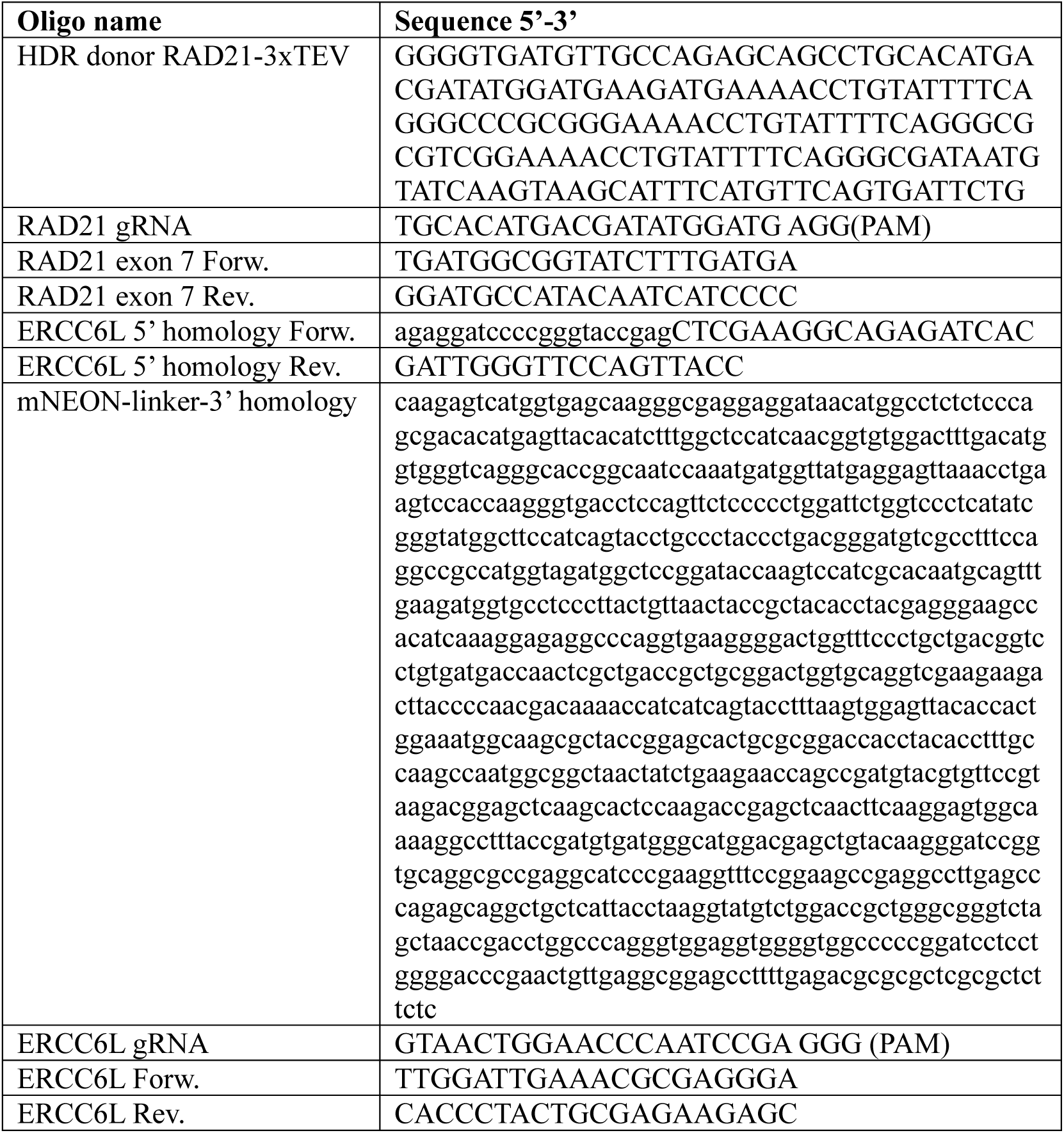
DNA oligos used in this study.

**Table S3.** Simulation parameters for buckling model.

| Category | Parameter | Value used |
| --- | --- | --- |
| *Springs* |  |  |
| Reduced extension | $u$ | $u = (L - L_0)/L_c$ |
| Contour-length scale | $L_c = \alpha L_0$ | $\alpha = 0.5$ |
| Spring constants by type | $k_{black}$ | $2.5 \times 10^3$ pN/μm |
| | $k_{yellow}$ | $4 \times 10^5$ pN/μm |
| | $k_{blue}$ | $4 \times 10^5$ pN/μm |
| Bending energy | B | 12 pN nm |
| *Distance sweep / numerics* |  |  |
| Displacement range | - | 0–0.4 μm |
| Increment | - | 0.005 μm |
| Energy minimizer | - | nonlinear conjugate gradient + backtracking line search |
| Max iterations per displacement step | - | 8000 |
| Stopping tolerance | - | Tolerance of $1 \times 10^{-7}$ per displacement step |
| Line search | - | initial step 1.0, shrink factor 0.8, min step $10^{-6}$ |
| *Measured outputs* |  |  |
| Extension | $\Delta x$ | mean x-displacement of pulled nodes |
| Stiffness measures | $k_{tan} \approx dF/d(\Delta x)$ | - |

**Movie S1.**

Chromosome attachment to Qtrap optical tweezers. A chromosome was first attached by a telomeric end to one of the beads. Then, the chromosome was flow-stretched to attach the other telomeric end to another bead. Then, while holding the chromosome taut, the remaining two beads were attached to the centromeric regions. Scale bar indicates 2 µm.

**Movie S2.**

Chromosome orientation is visible from spinning during attachment. Within the flow-cell, flow is applied from left to right. When the chromosome is attached to only one bead, this leads to spinning. This spinning can highlight the location of the centromere, indicated by the yellow arrow. Scale bar indicates 2 µm.

**Movie S3.**

Centromere stretching. A force of up to 250 pN was applied to the centromeres by moving the rightmost centromeric bead to the right and back. Scale bar indicates 2 µm.

**Movie S4.**

Centromere stretching after cohesin cleavage. Chromosomes were incubated with TEV protease to cleave cohesin prior to optical tweezers measurements (see Methods). Then once attached to the Qtrap optical tweezers, a force of up to 250 pN was applied to the centromeres by moving the rightmost centromeric bead to the right and back. Scale bar indicates 2 µm.

**Movie S5.**

YO-PRO-1 binding during centromere stretching. Chromosomes were first pre-treated with TEV protease, then trapped with the Qtrap optical tweezers. There they were incubated in a channel with YO-PRO-1 during stretching from 0-250 pN. Scale bar indicates 2 µm.

**Movie S6.**

TopIIα binding during centromere stretching. Centromeres were stretched up to 250 pN and back while inside a channel of the flow-cell containing TopIIα-A555. Scale bar indicates 2 µm.

